# Longitudinal changes in gut microbiota across reproductive states in wild baboons

**DOI:** 10.1101/2025.12.01.691429

**Authors:** Chelsea A. Southworth, Logan Barrios, Mauna R. Dasari, Laurence R. Gesquiere, Jack A. Gilbert, Susan C. Alberts, Jenny Tung, Elizabeth A. Archie

## Abstract

**Background:** In humans and other mammals, female reproduction is linked to extensive changes in physiology, immunity, hormones, and behavior. These changes likely shape, and may be shaped by, the composition of gut microbial communities. Characterizing the dynamics of gut microbial change across reproductive states, including its relationship to female physiology, is important for understanding how the gut microbiota influences female and offspring health.

**Results:** Here we characterize longitudinal changes in gut microbiota across reproduction by combining 16S rRNA gene sequencing data from 4,462 stool samples (spanning 14 years of sample collection) with life history data on multiple reproductive events in 169 female baboons. These baboons were members of a well-studied, natural baboon population in Kenya where reproductive state (ovarian cycling, pregnancy, and postpartum amenorrhea) is tracked daily and microbiota data could be paired with measurements of fecal-derived estrogen, progesterone, and glucocorticoid levels. We found extensive changes in baboon gut microbiota as females transitioned between reproductive states. Pregnancy was linked to distinct patterns of ASV richness, community composition, and taxonomic abundances compared to postpartum amenorrhea and ovarian cycling. The most dramatic shifts occurred as females transitioned from the first to second trimester of pregnancy, with altered abundances of taxa that have been linked, in humans and model systems, to host immunity, weight gain, or hormone levels. Host identity was consistently the strongest predictor of gut microbiota composition across states, and this individual signature was strongest during pregnancy. Estrogen and progesterone levels had robust associations with the gut microbiota overall, but the microbial taxa involved in these associations were reproductive state-dependent. Glucocorticoid concentrations were not a major predictor of gut microbiota composition in any state.

**Conclusions:** Together, our results support the idea that gut microbiota contribute to the complex physiological changes necessary during pregnancy, but that microbial changes during pregnancy are somewhat unique to each female. Variation in steroid hormones is associated with some, but not all, of these relationships, emphasizing the importance of considering steroid hormone levels in studies of gut microbiota variation. Our results motivate future work on how gut microbiota contribute to reproductive outcomes, including both maternal and offspring health.

## INTRODUCTION

Mammalian gut microbiota are highly complex, personalized microbial communities whose dynamics both reflect their host’s physical state and can have direct effects on host health [1, 2]. Changes in female reproductive state may be some of the most important drivers of these dynamics, but current evidence for these effects remains largely cross-sectional or limited to lab and domestic animals [3–6]. As female mammals transition from conceiving, to gestating, to caring for offspring, they experience changes in physiology, immunity, and behavior that could alter gut microbial composition, particularly during pregnancy [6–14]. Females’ social relationships also change across reproduction, and changes in the rate of social interaction could also change microbial exposures [8, 13, 15–18]. Gut bacteria may also play a causal role in some of these changes, influencing hormone levels, immunity, metabolism, and behavior [19–23]. For instance, experiments that transplanted microbiota from pregnant women to germ free mice implicate gut bacteria in weight gain during pregnancy [24], and certain gut microbes play a key role in shaping circulating estrogen levels [22, 25]. Understanding gut microbial dynamics across female reproduction is therefore relevant to maternal health, as well as to reproductive outcomes in humans and other animals, including preeclampsia, preterm birth, litter size, and the likelihood of entering estrus [26–29].

Despite multiple important roles for the gut microbiota in female reproduction, studies that trace longitudinal changes in female gut microbial communities as females transition between reproductive states are lacking. Such data are necessary to understand how gut microbial dynamics align with female reproduction, which aspects of these dynamics are shared across hosts, and which are unique to individual hosts [30, 31]. However, collecting longitudinal data on gut microbial composition is challenging: in humans, collecting repeated samples from the same individual over time and across reproductive states is difficult, while in most other mammals, one or more stages of female reproduction are cryptic, including ovulation and early pregnancy [32, 33]. This limitation makes it difficult to assign a given microbiota sample to a specific reproductive state.

Given that changes in steroid hormone physiology (e.g., estrogen, progesterone, glucocorticoids) are intimately involved in driving changes in mammalian reproductive state, studies of the reproductive dynamics of the gut microbiota would also ideally take hormonal changes into account. However, data on hormone levels and gut microbial composition are rarely paired, especially in longitudinal study designs [34, 35]. Where paired data are available, studies have typically focused on a narrow band of women’s lives (e.g., menopause) or hormonal disorders that disrupt normal reproduction [36–38]. Nevertheless, existing work points to importance of estrogen, progesterone, and glucocorticoids in altering gut microbiota [25, 39–43]. Indeed, administering estrogen and progesterone to ovariectomized mice maintains the gut microbial diversity found in control mice, while diversity decreases in non-hormone treated ovariectomized individuals [41]. Further, gut microbes directly contribute to hormone production: some gut microbes produce enzymes that deconjugate estrogen in the gut, returning it to circulation [25], and administering butyric acid, a short chain fatty acid that is a major product of Bacillota (Firmicutes) taxa, to porcine granulosa cells stimulates cellular secretion of both estrogen and progesterone in pigs [40]. However, while these studies indicate that hormone levels can be mechanistically linked to the composition and function of gut microbiota, whether steroid hormone levels and the gut microbiota vary across female reproduction in natural populations remains unclear. To our knowledge, only one study has connected hormones and gut microbiota longitudinally during normal reproductive transitions in a natural mammal population [44]. Mallott *et al.* (2020) tested relationships between the gut microbiota, fecal estrogen, and fecal progesterone using 68 fecal samples across cycling, pregnancy, and lactation from 10 wild female Phayre’s leaf monkeys. This study suggested that progesterone might play a role in linking reproductive state to microbial diversity and composition, but the small sample size made it challenging to detect associations between individual taxa and either reproductive state or hormone levels [44].

Here we address some of these gaps using a 14-year longitudinal dataset of 16S rRNA-based gut microbial profiles from 169 wild female baboons in Kenya (n=4,462 fecal samples; median samples per host=21; range=1-100 samples; a sub-set of a previously published data set [45, 46]). Each female was sampled during a median of 3 “reproductive events” (i.e., all the reproductive states and phases leading to the birth of a given offspring; range=1-10 reproductive events), with a median of 6 samples per reproductive event per female (range=1-46). We tracked both the female baboons and their gut microbiota as the females naturally transited between reproductive states [47] (Fig. 1). Baboons are a valuable reproductive model for human reproduction because, like humans, they breed year-round and have a 30-40 day ovarian cycle. However, unlike humans and most other mammals, they exhibit readily observable visual signals that allow accurate, non-invasive identification of the follicular and luteal phases of ovarian cycling, menstruation, pregnancy, and postpartum amenorrhea (PPA) [48–52] (Fig. 1A and Methods). Transitions between these reproductive states occur alongside well-characterized shifts in steroid hormones, including an estrogen spike before ovulation, increased progesterone in the luteal phase, and sharp increases in both hormones during the first trimester of pregnancy [49, 53–55] (Fig. 1A).

**Figure 1.**
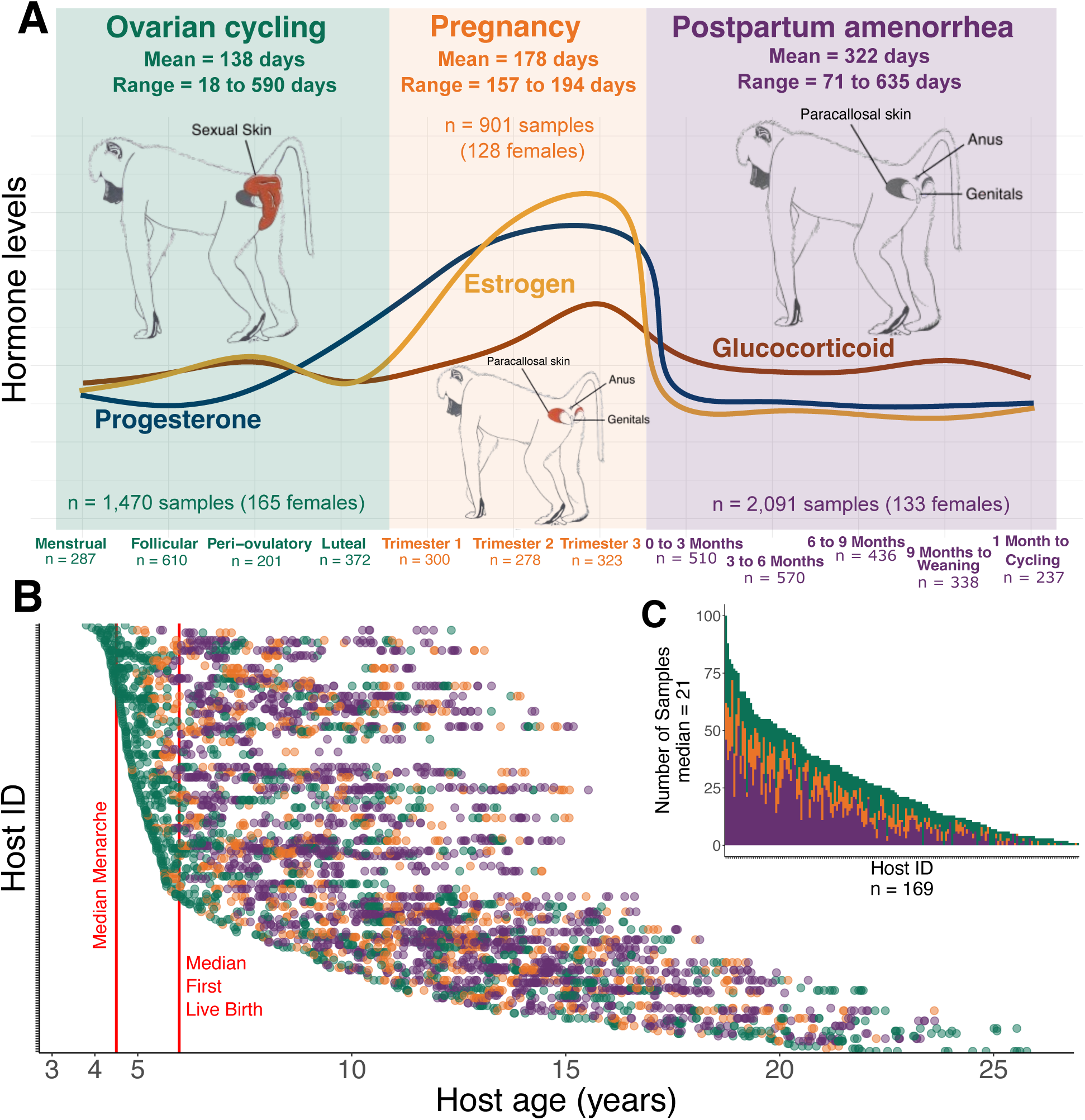
Baboon reproductive states and sampling design for this study. (A) Schematic showing reproductive states in baboons (ovarian cycling, pregnancy, and postpartum amenorrhea), mean lengths of each state (see Methods for description), and relative changes in estrogen, progesterone, and glucocorticoid hormones as female baboons transition between reproductive states. A full reproductive event was on average 638 days (range 333 to 1,084 days). Drawings within each state illustrate the external signals of that state in female baboons. Number of microbiota samples and female hosts are shown within each state. (B) Gut microbiota were characterized using 16S rRNA gene sequencing in 4,462 fecal samples collected longitudinally from each female host. The y-axis shows each female host, and each horizontal row of points represents the fecal samples collected for that female as a function of her age in years. Points are colored by reproductive state following the color scheme in (A). Median ages at menarche (4.5 years) and first live birth (5.97 years) are shown with red lines. (C) Number of samples collected (range=1-100, median=21) for each of the 169 female baboons in our dataset, colored by reproductive state.

Leveraging near-daily reproductive state data from the Amboseli population, our primary objective was to trace gut microbial changes as female baboons transitioned between ovarian cycling, pregnancy, and PPA. Our goal was to describe, in finer detail than available to date, both how and when the microbiota change across phases of female reproduction. We first tested for differences in microbiota between these states, controlling for known predictors of gut microbiota in this population, including season, weather, host diet, social group membership, and female age [45, 46, 56–60]. Based on prior research [11, 24, 61], we expected that gut microbiota would be most distinct during pregnancy because of this state’s profound changes in immunity, metabolism, and hormone levels [7, 10, 62]. We also tested how gut microbiota change within each reproductive state, but we did not have strong expectations about the patterns we would observe.

Next, we tested whether female gut microbiota exhibit reproductive personalization. It is well known that gut microbial communities are highly personalized (i.e., individual identity is often a key driver of gut microbiota composition) [46, 59, 63–67]. However, it is not known if this personalization is stronger during some reproductive phases than others or if individual females have particular microbial communities they return to in subsequent reproductive events across their lifetime. Because of the strong and predictable physiological changes during pregnancy, we expected that gut microbial composition would be the least personalized during pregnancy compared to ovarian cycling and PPA.

Finally, we tested how steroid hormone levels were associated with gut microbiota during each reproductive state, leveraging a unique feature of this data set: data on the concentration of fecal metabolites of estrogen, progesterone, and glucocorticoids measured in all 4,462 fecal samples [53, 68–71]. Based on previous research, we expected positive relationships between estrogen/progesterone levels and gut microbial alpha diversity, and a negative relationship between glucocorticoid concentrations and alpha diversity [36, 37, 72, 73]. Together, our results provide insight into gut microbial dynamics as a function of reproductive state, personalization, and hormones, highlighting the role of individualized host responses and host hormones in shaping this relationship. Our results will guide future research into the mechanistic basis and cross-mammalian consistency of these relationships.

## METHODS

### Study population and subjects

The female baboons we studied were the subjects of individual-based research by the Amboseli Baboon Research Project (ABRP) in Kenya. Baboons in this population are admixed between the yellow baboon (*Papio cynocephalus*) and the anubis baboon (*P. anubis*), with majority yellow baboon ancestry [74–76]. Prior research has found no link between host genetic ancestry and gut microbial composition in this system [77]. Since 1971, the ABRP has been collecting continuous observations of the baboons’ demography, behavior, and environment [47, 78]. The baboons are identified individually by experienced observers who collect data on each baboon social group 2 to 4 times per week; during the study period represented here, subjects could have lived in one or more of 10 different social groups [78]. Our study subjects were adult female baboons, i.e., females that had attained menarche. In Amboseli, menarche occurs at a median age of 4.50 years of age; the median age at first live birth is 5.97 years, and females continue reproductive activity into old age without evidence for systematic menopause [79, 80]. Data on female reproductive states and events (e.g., changes in ovarian cycle state, miscarriage, conception etc.) are collected from the study subjects on a near-daily basis, allowing for a fine-grained assignment of each subject to a reproductive state on each day during the study.

This research was approved by the IACUCs at Duke University and the University of Notre Dame and the Ethics Council of the Max Planck Society. It adhered to all the laws and guidelines of Kenya.

### Fecal sample collection, DNA extraction, and 16S data generation

#### Sample collection

The 4,462 gut microbial profiles in this analysis were a subset of 17,277 profiles that were previously described in [45, 46] (Fig. 1B and 1C). These 4,462 samples were collected from 169 individual adult females. The same fecal samples were also used to assay the concentrations of estrogen, progesterone, and glucocorticoid metabolites, providing paired microbiota-hormone data [70, 81, 82] (see below). Fecal samples were collected, stored, and extracted as described previously [70, 81, 82]; see Supplemental Methods.

#### DNA extraction, sequencing, and post-sequencing processing

DNA was extracted using the MoBio and QIAGEN PowerSoil kit for 96-well plates [83] with modifications described in [45, 46]; see Supplemental Methods. Sequences were processed using the Illumina demultiplexing protocol and DADA2 pipeline [45, 46, 84]; see Supplemental Methods. Taxonomic identification was performed using the *DECIPHER* package [85] against the Silva reference database SILVA_SSU_r138_2_2024.RData [86]. We removed samples that had DNA extraction concentrations <4X the blank on that sample’s plate [45]. Prior to analysis, we also excluded any amplicon sequence variants (ASVs) found in fewer than 5% of the 4,462 samples used in this analysis.

#### Quantifying gut microbial features

To measure within-sample alpha diversity, we calculated the ASV richness and Shannon diversity for each sample using the *estimate_richness* function from the *phyloseq* package in R [87, 88]. We also calculated Faith’s phylogenetic diversity using the *pd* function in the *picante* package [89, 90]. For these calculations, phylogenetic relationships between ASVs were estimated by first performing a multiple sequence alignment with the *DECIPHER* package [85]. A Maximum Likelihood tree was then constructed with the *phangorn* and *ape* packages, utilizing a Neighbor-Joining starting tree and the General Time Reversible model with Gamma-distributed rate variation and a proportion of invariable sites (GTR+G+I) [91, 92]. The final tree was midpoint rooted using the *phytools* package.

To measure differences in microbial community composition between samples, we centered log-ratio (CLR) transformed ASV relative abundances with the *microbiome* package’s *transform* function [93]. We then constructed an Aitchison distance [94] matrix using the *microbiome* package’s *dist* function [93]. This distance metric is the Euclidean distance between CLR-transformed samples and is an appropriate metric for handling the compositionality of microbial data; it is also robust to sub-setting samples [46, 95, 96]. To account for phylogenetic relationships among ASVs, we also calculated both weighted and unweighted UniFrac distances with the *distance* function in *phyloseq* [87]. These distance matrices used the same phylogenetic trees described above. Finally, we used the CLR-transformed relative abundances of individual microbial taxa to measure their differential abundance between samples. We focused on taxa that were commonly found across samples, specifically 401 ASVs, 50 families, and 14 phyla present in 20% or more of the 4,462 samples.

### Measuring predictors of the gut microbiota across reproductive states

To understand how the microbiota changes as a function of reproductive state, and whether and to what degree individual identity and hormone profiles were associated with these changes, our primary variables of interest were: (i) host reproductive state and phase; (ii) the identity of the reproductive event; and (iii) the concentrations of estrogen, progesterone, and glucocorticoid metabolites in a given sample. Below we describe how these variables were measured, followed by a description of the other variables we included in our analyses.

#### Reproductive variables

*(i) Reproductive state and phase.* Following menarche, female baboons move through three reproductive states: ovarian cycling, pregnancy, and postpartum amenorrhea (PPA), all of which are easily distinguishable by external signals [49, 51, 97] (Fig. 1A). Each fecal sample for microbial analysis was assigned to one of these states, with further sub-categories within each state described below.

##### Ovarian cycle data set

An ovarian cycle consists of four phases: (1) the follicular phase, during which estrogen concentrations rise and the sexual skin begins to swell; (2) the peri-ovulatory phase, i.e., the five-day ovulation window when estrogen peaks and the sexual swelling reaches its maximum size and turgescence; (3) the luteal phase, which is marked by the deturgescence of the sexual skin and a concurrent decrease in estrogen and increase in progesterone; and (4) the anestrus/menstrual phase, where the sexual skin becomes flat and menstrual blood may be observed if a female does not conceive [49, 68, 69, 97]. The mean duration of ovarian cycling from cycle resumption to conception for females in the Amboseli population is 138 days (SD=82, range=18-590 days), which equates to 6-8 cycles to conception [68, 69].

For samples in this ovarian cycle data set (hereafter cycling samples), we only included normal ovarian cycles where the swelling length (i.e., the combined follicular phase and 5-day periovulatory phase) was between 10 and 39 days long. This criterion excluded 4% of the total samples in the pre-filtered data set, including unusually long and short cycles that occur in especially young or old females. Our final data set for cycling females therefore included a total of 1,470 fecal samples from 165 female baboons collected during 407 distinct periods of ovarian cycling.

##### Pregnancy data set

Pregnancy is identified by the cessation of sexual cycling (i.e. the absence of sexual skin swellings and menstruation) and a transition of the paracallosal skin (PCS) from gray to pink during the first several months following conception (the baboon “pregnancy sign”) [53, 97]. While this visual assessment correctly identifies 97% of endocrinologically-confirmed pregnancies, we are likely to miss pregnancies where fetal loss occurs early in pregnancy [53]. In Amboseli, the average gestation for a live birth is 178 days (SD=6, range=157-194), which we divided into three trimesters of 60 days each [53, 69]. We only included samples from pregnancies that resulted in a live birth, resulting in a total of 901 fecal samples collected during 359 distinct pregnancies from 128 females.

##### Post-partum amenorrhea data set

Once a female gives birth, her PCS typically turns from pink back to dark gray (although females in our population and in other baboon species can retain some permanent pink after a pregnancy, and the amount of ‘permanent pink’ skin can increase with each parturition) [53, 97–99]. Females then enter a period of post-partum amenorrhea (PPA) during which their PCS remains flat. After giving birth to an infant, females whose infants survived their first year of life spent an average of 322 days (SD=87, range=71-635 days) in PPA before cycling resumption [69].

We divided PPA into five periods based on the energetic demands of the infant and physiology of the mother. Each fecal sample for microbiota and hormone analysis was assigned to one of these five periods: (1) 0-3 months postpartum, when infants are completely reliant on the mother for nutrition and transportation; we excluded the first seven days postpartum due to the high hormone levels that persist immediately after a recent pregnancy; (2) 3-6 months postpartum, when infants begin to ingest food other than milk but still heavily rely on maternal carrying; (3) 6-9 months postpartum, when infants became more independent but still ingest milk and are still carried during periods of rapid movement despite their larger size; (4) 9 months postpartum to infant weaning (∼70 weeks postpartum), when infants are largely eating and traveling independently; and (5) one month prior to cycling resumption, when female energetic condition begins to improve after the metabolic demands of lactation [69, 70, 100–102]. Note that this fifth category overrode the others: e.g., if a female resumed cycling in month 8, a fecal sample in month 8 was assigned to the 30 days prior to resumption rather than the 6-9 months postpartum period.

We excluded samples from PPAs than lasted more than 496 days (>2 SD from the mean of 337.5 days in our pre-filtered data set). Likewise, because we excluded pregnancy samples if a miscarriage occurred, we also excluded samples from the (typically brief) PPA periods that followed miscarriages. If a female gave birth to a live infant that died while the female was in PPA, samples from that PPA were included if the sample was collected prior to the infant death or if the sample was collected after the infant death but within 30 days of the female’s cycling resumption. Samples collected after the infant death but >30 days before cycling resumption were excluded, as they did not fit into any of the PPA phase categories as we have defined them above. The final PPA samples therefore included a total of 2,091 fecal samples from 133 females collected during 394 periods of PPA.

*(ii) Reproductive event.* We used the term “unique reproductive event” (n=551) to refer to the complete set of reproductive states a female experienced starting with menarche or cycle resumption (after the birth or fetal loss of a previous infant), continuing through the conception and birth of an offspring and the subsequent lactation period, and ending when the female resumed cycling again in preparation for the next conception [69] (Fig. 1). A full reproductive event for a female whose infant survived to one year was on average 638 days long (SD=116, range=333-1084 days[69]). When a female resumed cycling after a conception, whether because the fetus or infant died or because the mother resumed cycling once the infant survived to near-independence, the first day of cycling marked the beginning of the next reproductive event. Because the baboons were observed on a near-daily basis, the length of each unique reproductive event in our data set was accurate to within a few days’ error.
*(iii) Steroid hormone concentrations.* Steroid hormone concentrations of fecal estrogen (fE), progesterone (fP), and glucocorticoid (fGC) metabolites were measured in all fecal samples via I-125 radioimmunoassay, using a well-established protocol described in [70, 81, 82]. Hormone metabolites were expressed as nanograms of hormone per gram of lyophilized, sifted fecal sample.

#### Non-reproductive variables known to predict microbiota composition

In addition to the three reproductive variables discussed above (reproductive state/phase, reproductive event, and steroid hormone concentrations), we also included as predictors in our models several other variables known to predict gut microbial composition in our study population [45, 46, 56–60]. The results for these variables were largely consistent with prior studies [45, 46, 56–60]; hence, in the results, we focused solely on outcomes for reproductive state and hormone concentrations. These non-reproductive variables included: (iv) host traits such as host identity, age, and social group membership, (v) measures of host diet in the 30 days before sample collection, (vi) rainfall in the 30 days before sample collection, and (vii) technical aspects related to 16S rRNA profile data generation. In exploratory analyses, we also tested for effects of season (wet/dry) and year, but these were excluded due to strong correlations with diet and rainfall. Although gut microbiota phenotypes are heritable in our study population [45], we did not control for genotype because estimated heritability values are quite small and we sought to avoid overparameterizing our models, which are already complex.

*(iv) Host traits (individual identity, age, social group membership).* All baboons (n=169) were individually recognizable on sight by ABRP observers and born into continuously observed study groups. As a result, age for most baboons (n=158) was known to within a few days’ error. For eleven females, age was estimated to within six months’ error, i.e., the birth date was estimated with an error of ± three months. Group membership was recorded from near-daily censuses of all group members (n=10 social groups) [78]. Group membership and age are known to predict microbiota composition in the Amboseli baboons and were included in our analyses as controls [45, 46, 56–58, 60].
*(v) Diet.* Host diet composition was measured as described in [45, 46] and included in our analyses to control for known diet-dependent variation in microbiota composition. Briefly, data on diet were collected during 10-minute focal animal samples of adult females and juveniles [78, 103]. When the focal animal was observed eating, the food type and part was recorded. These food types were grouped into 14 categories (e.g., grass blades, grass seed heads, blossoms, etc.). Because feeding data collected within each focal animal sample are autocorrelated, we randomly generated 1,000 subsets of 1 feeding value per focal sample. An individual’s diet was estimated using feeding observations from all focal samples in the individual’s social group during the 30 days prior to fecal sample collection. Each fecal sample was thus assigned a specific set of values representing the proportion of time spent feeding on each of the 14 food categories. Several studies support the use of social group-level diet data to represent individual-level diet in the Amboseli baboons [104–109]. To ensure that diet composition data were reliable, we only included fecal samples for microbial analysis when at least 15 focal samples were collected in an individual’s social group in the 30 days prior to when the fecal sample was collected [45]. We calculated the dietary Shannon’s diversity for each fecal sample using the *diversity* function in the R *vegan* package to quantify diversity and evenness of the diet composition associated with each fecal sample [88, 110]. To measure compositional variation in diet between fecal samples, we conducted a principal components analysis (PCA) on the diet matrix of the 14 food categories using the base R *prcomp* function.
*(vi) Rainfall.* Rainfall is highly seasonal in the Amboseli ecosystem and has profound effects on food availability for the baboons [78, 104, 106, 111]. Hence, rainfall was included as a predictor variable in all our models to control for known rainfall-dependent variation in microbial composition [45, 46, 56, 57, 60]. Rainfall data were collected using a rain gauge at the ABRP camp and represented as the sum of total rainfall during the 30 days prior to sample collection [78].
*(vii) Technical variables (sequencing depth, plate ID).* Sample sequencing depth (read count) was included to control for apparent differences in microbial diversity and abundance due to technical variation and was quantified using the *DADA2* pipeline [84] (range=1,028-477,241). Samples were loaded onto 166 different extraction plates and plate identity was included as a predictor variable to control for batch effects.

### Statistical analyses

All statistical analyses were performed in the R statistical environment (version 4.2.0 [88]).

#### Aim 1: Testing gut microbiota changes between and within reproductive states

To examine the relationship between microbial alpha diversity, reproductive state (cycling, pregnant, or in PPA), and phases within states (pregnancy trimesters, phases of cycling or stages of PPA), we ran a series of linear mixed effects models using the *lme4* and *lmerTest* packages [112, 113]. The response variables in these models were our three measures of alpha diversity: ASV richness, Shannon diversity, and Faith’s phylogenetic diversity. The key fixed effect was (1) female reproductive state or phase (e.g., cycling, pregnant, or in PPA, or pregnancy trimester for models sub-set to pregnant females). To control for other known sources of microbial composition we also included fixed effects of: (2) host age in years; (3) total rainfall in the 30 days prior to sample collection in mm; (4) the sequencing depth of the sample; (5) the collection date of the sample as a running count from the start of the sample collection period; (6) dietary Shannon’s diversity calculated using the *vegan* package [110]; and (7-11) the first five principal components of the dietary PCA, which explained a cumulative 55.36% proportion of the variance in diet between samples (we selected the minimum number of PCs that explained >50% of the variance in diet composition to avoid overparameterization). To facilitate comparison of effect sizes, all fixed effects were standardized to mean zero and variance of 1 across the entire data set. We also modeled random effects of host baboon identity, the host’s social group at time of collection, and the sample extraction plate as a technical control.

To test for differences in microbial community composition between hosts in different reproductive states and phases, we ran ANOVAs using sequential marginal tests (999 permutations) on redundancy analysis (RDA) models predicting variance in the CLR-transformed microbial abundance matrix using the *vegan* package [110]. To test for differences in phylogenetically informed measures of community composition, we calculated weighted and unweighted UniFrac distance matrices and ran ANOVAS using sequential marginal tests (999 permutations) on distance-based redundancy analysis (dbRDA) models predicting variance in these matrices. Individual identity was included as a stratum in the ANOVA, and the same variables used in the taxonomic richness model were included as fixed effects in the RDA and dbRDAs (host reproductive phase, host age, rainfall, sample read count, sample collection date, measures of host diet, host social group, and sample extraction plate). We chose this method because the variance in the distributions of several key variables was significantly different (e.g., reproductive states: weighted UniFrac matrix ANOVA F=5.69, p=0.003, unweighted UniFrac matrix ANOVA F=7.35, p<0.001; trimesters of pregnancy: unweighted UniFrac matrix ANOVA F=6.48, p=0.002; phases of PPA: CLR-transformed abundance matrix ANOVA F=4.23, p=0.002, unweighted UniFrac matrix ANOVA F=2.44, p=0.045; from *betadisper* in *vegan*), violating a key PERMANOVA assumption.

To test the effect of reproductive state and phase on the abundances of individual microbial taxa, we ran linear mixed effects models on the CLR-transformed relative abundances of the 401 ASVs, 50 families, and 14 phyla that were present in 20% or more of samples. The model structure was the same as the linear models of alpha diversity. For each set of analyses (across states, within pregnancy, within PPA, and within cycling), we controlled for false discovery using the Benjamini-Hochberg correction. Adjustments were performed independently for each microbial taxonomic level and for all pairwise comparisons between values for each variable of interest (reproductive state or phase within state). For interpretability, we focus our presentation of the results on taxa that had relatively strong effect sizes (>0.4 or <-0.4, equivalent to a fold change ∼1.5 or −1.5) and a q-value <0.05. However, it is possible that changes below this threshold could be biologically important, and we therefore present results for all taxa in our supplementary tables.

#### Aim 2: Testing for gut microbiota personalization between and within reproductive states

To test the effect of individual identity and reproductive state on gut microbial beta diversity between reproductive states, we subset our data to the ten best-sampled hosts (range=67-100 samples per female). As described above, we ran ANOVAs (sequential marginal tests, 999 permutations) on RDAs of CLR-transformed abundances, and dbRDAs of weighted and unweighted UniFrac distance matrices for each female’s samples separately. Predictors included host reproductive state as well as most of the other host, environmental, and technical variables described in Aim 1. To avoid overfitting these smaller data sets, we simplified the model structure by removing the host social group and age variables as there was insufficient variation in either variable at the within-host level; host age also explained only a small portion of variation between samples.

To measure personalization within reproductive states, we subset the full data set to adult females who had at least 10 total samples with at least two samples from each reproductive state, and at least two samples that were from the same reproductive event (N=4,137 samples from 116 females and 452 reproductive events). We then used temporal autocorrelation analysis to reveal the extent to which longitudinal patterns of microbial change were specific to individual reproductive events, to individual females, or to the reproductive state in general (i.e., across females and reproductive events). To accomplish this goal, we first transformed the Aitchison, weighted UniFrac, and unweighted UniFrac distances between samples into Aitchison, weighted UniFrac, and unweighted UniFrac similarities bounded from 0 to 1, where a similarity of 1 indicates that two samples are identical and a similarity of 0 indicates that two samples are completely different [46]. We then tested similarities between three different categories of sample pairs in each reproductive state: (1) pairs from the same reproductive event in the same female, (2) pairs from different reproductive events in the same female, and (3) pairs from different females in the same social group during the same reproductive state.

For these categories (e.g., within female, same pregnancy; within female, different pregnancy; between pregnant females, same social group), we tested similarities between sample pairs as a function of lag times of reproductive-months. For example, two samples collected on day 40 and day 60 of pregnancy would be within one reproductive-month lag of each other, regardless of the calendar dates on which the samples were collected. Hence, for comparisons that fall in category 1 above, for pregnancy, the two samples would have been collected on the 40^th^ and 60^th^ day of the same pregnancy from the same female. For category 2, the two samples would have been collected on the 40^th^ day of one pregnancy and the 60^th^ day of a different pregnancy from the same female. Thus, they were most likely to be have been collected in different calendar years (e.g. one sample in March 2005 on the female’s 40^th^ day of one pregnancy and one sample in November 2010 on the female’s 60^th^ day of a different pregnancy). Category 1 comparisons were necessarily collected chronologically close in time to each other (a one reproductive-month lag is also one chronological-month lag) and category 2 comparisons were necessarily always further in chronological time to each other (a one reproductive-month lag is always longer than a one chronological-month lag, due to the time between different reproductive events in the same female). Category 3 comparisons, which contrast females living in the same social group, include samples of varying chronological-month lags relative to the reproductive-month lags.

Within each of the three comparison categories, we considered pairwise comparisons of samples collected with reproductive-month time lags ranging from 1 to 6 months. Reproductive-month time lags of one month correspond to samples collected within 0-30 days of each other. Time lags of two months correspond to a 30–60-day difference on the reproductive timeline, and so on, up to a maximum difference of 150-180 days (six reproductive-months). Because we consider a maximum of six months, the length of a typical successful pregnancy, this approach means that we track more pairwise comparisons for time lags of one month than for six months.

For all comparisons, we calculated the mean and upper and lower confidence intervals across samples in each reproductive state (considering pregnancy, cycling, and PPA) [114]. For example, we calculated the mean Aitchison weighted UniFrac, and unweighted UniFrac similarity and confidence intervals across pairs of pregnancy samples from the same pregnancy (category 1) at one, two, three, four, five, and six reproductive-month time lags, then for pairs of pregnancy samples from different pregnancies in the same female (category 2) across each of the six time lags, and finally for pairs of pregnancy samples from pregnancies in different females (category 3) across each of the six reproductive-month time lags. We repeated these calculations for cycling and PPA samples. To test the statistical significance of differences between similarity in means within each reproductive state at each reproductive-month time lag, we conducted pairwise t-tests with Bonferroni-corrected p-values [115].

#### Aim 3: Testing gut microbiota associations with steroid hormone estrogen, progesterone, and glucocorticoid metabolite concentrations

To examine the relationship between fE, fP, and fGC and microbial alpha diversity, we ran linear mixed effects models with the general structure described in Aim 1 (including host reproductive state or phase and host, environmental, and technical covariates). Here, we modeled ASV richness, Shannon diversity, and Faith’s phylogenetic diversity as the response variables, with the addition of fE, fP, and fGC metabolite concentrations as fixed effect predictors. For each hormone, its fecal concentration (ng/g dry fecal matter) was first natural log transformed and modeled as a function of days from collection to extraction and days from extraction to assay (log fecal hormone concentration ∼ days to extraction + days to assay) to control for known effects of time in storage [116]. We extracted the residuals from these models and mean-centered each set of residuals to zero for each hormone separately. While hormones are correlated both with each other and with reproductive states and phases (Fig. 1A), Variance Inflation Factor scores [117] were all <5, a conservative estimate for mathematically problematic multicollinearity. We modeled all samples from all reproductive states together, including interaction effects between each hormone and reproductive state to test if the effects of hormones on alpha diversity varied between reproductive states.

To test for a relationship between fE, fP, fGC, and beta diversity, we ran ANOVAs (sequential marginal tests, 999 permutations) on RDAs (CLR-transformed abundance matrix) and dbRDAs (weighted and unweighted UniFrac distance matrices) of all samples and within each reproductive state separately. As described in Aim 1, the RDA and dbRDA models included reproductive state or phase, fixed effects of host, environmental, and technical covariates, in addition to standardized fE, fP, and fGC concentrations corrected for technical effects of collection and extraction time in the models. As for our analyses of alpha diversity and steroid hormone metabolites, when modeling all samples together, we included interaction effects between each hormone and reproductive state to statistically test if the relationships between hormones on beta diversity varied between reproductive states. Individual identity was included as a stratum in the ANOVAs.

To test for a relationship between fE, fP, fGC, and the abundances of individual microbial taxa, we ran linear mixed effects models on CLR-transformed taxonomic abundances at the phylum, family, and ASV levels for taxa present in 20% or more of samples. False discovery rate adjustments with the Benjamini-Hochberg correction were performed independently for each set of analyses (across states, within pregnancy, within PPA, and within cycling) and each microbial taxonomic level. These models included standardized fE, fP, and fGC concentrations corrected for technical effects, in addition to the variables described in the methods for Aim 1 including host reproductive state or phase and measures of host, environmental, and technical covariates. We ran these models across all samples to test for generalized, consistent relationships between hormones and microbial taxa, and additionally ran these models within each reproductive state separately to test differences between states in hormone-taxon relationships.

## RESULTS

### 1. Gut microbial composition changes between reproductive states, and the pregnancy microbiota is distinct

After quality control and filtering, we detected a total of 898 microbial ASVs with a prevalence >5% across all 4,462 samples. On average, each sample had 240 ASVs (SD±79) with representatives from a range of microbial families, including members of Bifidobacteriaceae, Prevotellaceae, Lachnospiraceae, and several other families typically observed in primate gut microbiota. Each sample had an average Shannon diversity of 3.49 (SD±0.62) and an average Faith’s phylogenetic diversity of 51.63 (SD±8.96).

#### Gut microbiota change predictably between reproductive states

Baboon gut microbiota exhibited predictable changes in composition as females transitioned between reproductive states (Fig. 2, Fig. S1). Microbial diversity was highest in pregnant females, who harbored 13 more ASVs, on average, than cycling females, and 10 more ASVs than females in PPA, or about a 5% increase from the average sample’s ASV richness. While small, this effect exceeds that of host age, measures of host diet, and cumulative rainfall (Fig. 2A; Table S1). Both Shannon diversity and phylogenetic diversity were also higher in pregnant females than those in ovarian cycling (3.2% and 2.4% increase, respectively, from the mean diversity) or PPA (4% and 1.9% increase, respectively, from the mean diversity; Fig. S2; Table S1).

**Figure 2.**
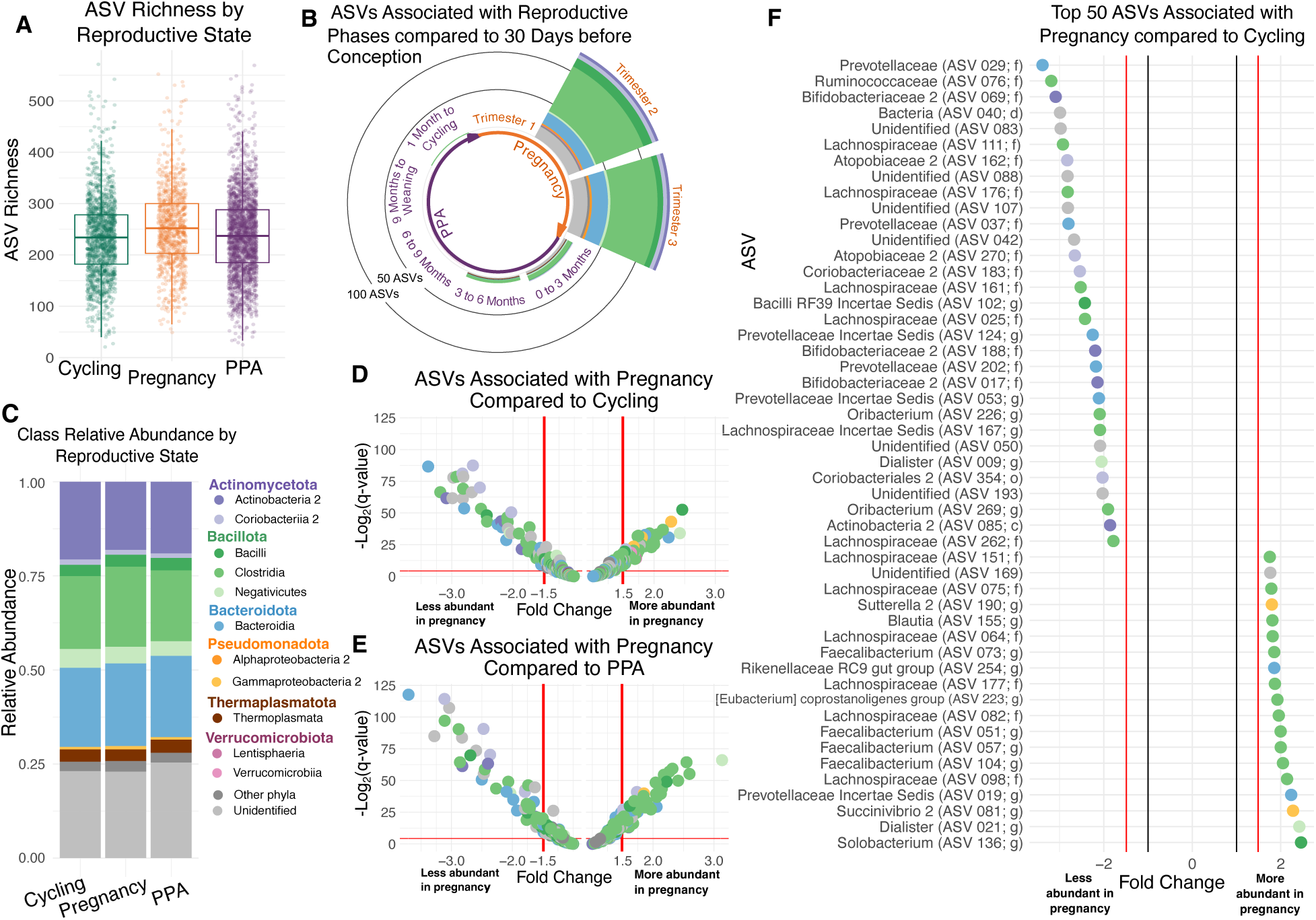
Gut microbiota change as females transition between reproductive states and are especially distinct during pregnancy. (A) ASV richness is higher during pregnancy than ovarian cycling (β=12.8, p<0.001) or PPA (β=10.0, p<0.001). Full model results are in Table S1. (B) Several ASVs change substantially in abundance (i.e., fold change β1.5 or ≤-1.5 and q<0.05) as females transition between reproductive states, especially members of the phyla Bacillota (Firmicutes) and Bacteroidota (Bacteroidetes). All changes in abundance are shown in reference to the 30 days prior to conception. Analyses were performed on the 401 ASVs present in 20% or more of samples. Model results used to generate this figure are in Table S3. (C) Average relative abundance of bacterial classes from one reproductive state to the next. “Unidentified” taxa are those not identified to the class level and “Other” taxa aggregates rare taxa identified to the class level but present in <40% of samples. (D and E) Volcano plots showing the effect of pregnancy compared to (D) ovarian cycling and (E) PPA on the abundances of the 401 ASVs. Each point represents an individual ASV and the color of each point represents the ASV’s assigned class. Points above the horizontal red line are statistically significant (q<0.05). Model results are in Table S4. (F) The 50 ASVs with the greatest differences in abundance in samples collected during pregnancy compared to ovarian cycling. Point color represents the assigned class of each ASV. The black vertical lines at −1 and 1 represent the minimum possible fold changes. Model results are in Table S4.

Changes in overall between-sample beta diversity between reproductive states were small (CLR abundance matrix R^2^=0.0146, p=0.001; weighted UniFrac matrix R^2^=0.0030, p=0.001; unweighted UniFrac matrix R^2^=0.0037, p=0.001; Fig. S3; Table S2). However, the number of microbial taxa that changed in abundance between reproductive states was relatively large—especially during pregnancy (Fig. 2B-2F). For instance, when females transitioned from cycling to pregnancy, they exhibited changes in the abundances of 92 ASVs (22.9% of 401 tested ASVs in >20% of samples), a majority of which were butyrate-producing Bacillota (Firmicutes) ASVs (Fig. 2D, 2F; Table S4). Pregnant females also had higher abundances of 5 families (vadinBE97, Succinivibrionaceae 2, Sutterellaceae 2, Oscillopsirales UCG-010, and an unidentified Burkholderiales 2 family) and the phylum Verrucomicrobiota (Lentisphaerae), as well as lower abundances of 3 families (an uncultured Bacteroidales family, an unidentified Coriobacteriales 2 family, and an unidentified Actinobacteria 2 family) compared to cycling females (Fig. 2D, 2F; Table S4). Furthermore, when females transitioned from pregnancy to PPA, they exhibited changes in the abundances of 97 ASVs (24.2% of 401 ASVs), which were also dominated by Bacillota ASVs (Fig. 2E; Table S4). 5 families also shifted in abundance between pregnancy and PPA (Succinivibrionaceae 2, Sutterellaceae 2, and Erysipelatoclostridiaceae lower in abundance; an unidentified Coriobacteriales 2 family and an uncultured Rhodospirillales 2 family higher in abundance), but no phyla changed in abundance (Fig. 2E; Table S4). By contrast, as females transitioned from PPA to ovarian cycling, they exhibited changes in the abundances of just 7 taxa, including lower abundances of the family vadinBE97 and changes in the abundances of 6 ASVs (1.5% of 401 ASVs; Fig. S4; Table S4).

#### Gut microbiota shifts during pregnancy emerge in the second and third trimesters

Baboon gut microbiota also changed within pregnancy. The first trimester was similar to the cycling microbiota, while the larger microbial differences that characterized pregnancy as a whole emerged during trimesters 2 and 3 (Fig. 2B, Fig. 3; Table S5-7). Alpha diversity increased slightly over the course of pregnancy, with trimesters 2 and 3 showing higher ASV richness, Shannon diversity, and phylogenetic diversity than trimester 1 (Fig. 3A, Fig. S5; Table S5; trimesters 2 and 3 did not differ for any of our three measures of alpha diversity). We observed small changes in microbial beta diversity between trimesters (CLR abundance matrix R^2^=0.0064, p=0.001; weighted UniFrac matrix R^2^=0.0040, p=0.039; unweighted UniFrac matrix R^2^=0.0082, p=0.001; Table S6). In terms of individual taxa, 129 ASVs changed in abundance when females in the 2^nd^ trimester of pregnancy are compared to females in the 30 days prior to conception (Fig. 2B; Table S3). 8 families and 72 ASVs (18.0% of 401 ASVs), particularly Bacillota ASVs, differed in relative abundance both in the pairwise comparisons of trimester 1 and 2, and when comparing trimester 1 and 3 (Fig. 3B, 3D-E; Table S7). An additional 31 ASVs, 6 families, and 2 phyla varied in abundance between trimesters 1 and 2 (but not 1 and 3), while an additional 27 ASVs and 1 family differed in abundance between trimesters 1 and 3 (but not 1 and 2) (Fig. 3B, 3D-E; Table S7). These families included several members of phylum Bacillota, such as Lactobacillaceae, Monoglobaceae, Erysipelatoclostridiaceae, Selenomonadaceae, and Oscillospirales UCG-010 (Table S7). In contrast, only one ASV, unidentified at the domain level, differed in abundance between trimesters 2 and 3 (Fig. 3C, 3F; Table S7). These patterns indicate that major pregnancy microbial shifts begin early in the second trimester and stabilize by the third. See Table S7 for details on specific taxa.

**Figure 3.**
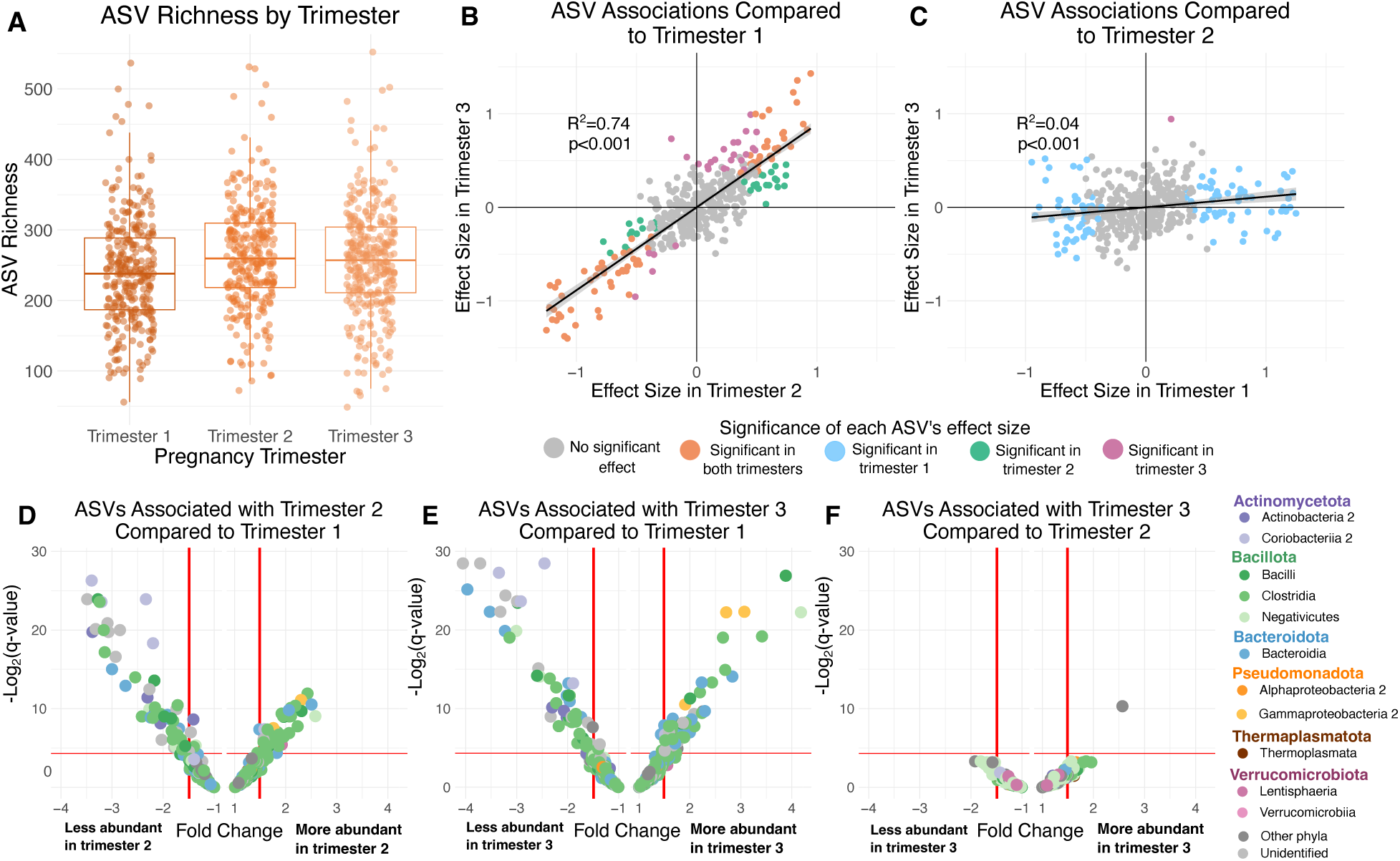
Gut microbiota change across pregnancy, with the “pregnancy microbiota” emerging during trimesters 2 and 3. (A) Trimesters 2 and 3 have higher ASV richness than trimester 1 (β=19.1, p<0.001 in trimester 2, β=12.1, p=0.007 in trimester 3; see Fig. S5 for Shannon and phylogenetic diversity; Table S5). (B and C) Scatterplots showing the correlation between effect sizes for individual ASV abundances in (B) trimester 2 and trimester 3 compared to reference state trimester 1 and (C) trimester 1 and trimester 3 compared to reference state trimester 2. Each point represents one of the 401 ASVs present in 20% or more of samples, colored by the trimester(s) where there is a significant (q<0.05) difference in CLR-transformed relative abundance from the reference state. R^2^ and p-values are from linear models on the points, with linear model best fit lines shown on the plots. (D, E and F) Volcano plots showing the differences between (D) trimester 2 compared to trimester 1, (E) trimester 3 compared to trimester 1, and (F) trimester 3 compared to trimester 2 on the abundances of the 401 ASVs. Each point represents an individual ASV and the color of each point represents the assigned class of the ASV. Points above the horizontal red line are statistically significant (q<0.05). Model results used to generate figures B-F are in Table S7.

#### Early postpartum amenorrhea (PPA) differs microbially from later PPA

Baboon gut microbiota in early PPA were similar to those at the end of pregnancy, with the microbial features that characterized PPA emerging after 3 months postpartum. For example, in an extension of the elevated alpha diversity in trimesters 2 and 3 (Fig. 3A), ASV richness and Shannon diversity were significantly higher in the first 3 months of PPA compared to females in all later phases of PPA (ASV richness: β=-12.72 to −6.63, p≤0.026; Shannon diversity: β=-0.18 to −0.085, p≤0.016; Fig S6A-B; Table S5). Phylogenetic diversity did not significantly differ between the first 3 months of PPA and months 3-6 (β=-0.61, p=0.095), but it was higher during the first three months of PPA compared to the later three phases after 6 months postpartum (β=-1.55 to −1.15, p≤0.017; Fig. S6C; Table S5).

Small changes in beta diversity also occurred as females transitioned between periods of PPA (CLR abundance matrix R^2^=0.0030, p=0.001; weighted UniFrac matrix R^2^=0.067, p=0.001; unweighted UniFrac matrix R^2^=0.0041, p=0.001; Table S6). In terms of individual taxa, 64 ASVs (16% of 401 ASVs), 21 families, and 2 phyla changed in relative abundance between the first three months of PPA and at least one other period of PPA (Fig. S6D-G; Table S8). Many of these ASVs belonged to phylum Bacillota, and among the associated families were nine Bacillota, including Acidaminococcaceae and Streptococcaceae, as well as Lactobacillaceae, Monoglobaceae, Selenomonadaceae, and Oscillospirales UCG-010 which also differed in abundance between pregnancy trimesters (Table S7-8). See Table S8 for details on specific taxa.

#### Gut microbiota do not show strong changes across ovarian cycle phases

Despite the fact that baboon female physiology, behavior, and vaginal microbiota change across the 4 phases of the ovarian cycle [18, 68, 118, 119], we found very few changes in baboon gut microbiota. No alpha diversity metrics changed as a function of ovarian cycle phase (Fig. S7A-C; Table S5), nor did beta diversity (p≥0.1.78 for all distance matrices; Table S6). Only one taxon varied between ovarian cycle phases at the q<0.05 and estimate <-0.40 or >0.40 cutoffs: the phylum Thermodesulfobacteriota was more abundant in menstruating females than those in peri-ovulation (Fig. S7D-F; see Table S9 for details on all taxa).

### 2. Changes in the microbiota with reproductive state are individualized, especially during pregnancy

Many studies have shown that gut microbial communities are highly personalized to individual hosts, including in the Amboseli baboons [46, 59, 67]. Thus, we wondered if the effects of reproductive state on gut microbial composition were stronger when inferred based on samples collected from the same female host. In support of this possibility, when we subset our data to the 10 best-sampled females (those with ≥65 samples; range=67-100 samples), the percent variance in community composition attributable to reproductive state was 2 to 3.6 times higher (CLR-transformed abundance matrix), 9 to 29 times higher (weighted UniFrac matrix), and 6.5-19.5 times higher (unweighted UniFrac matrix) for samples from a single host than for all hosts combined (Fig. S8; Table S10). The percent variance in the CLR-transformed abundance matrix explained by reproductive state was statistically significant within all hosts and ranged from 2.8% to 5.4% (Fig. S8; Table S10). Reproductive state explained significant variation in the weighted UniFrac matrix in 6 of 10 hosts and in the unweighted UniFrac matrix in 8 of 10 hosts, with percent variance ranging from 3.4% to 8.8% and 3.7% to 7.2% for each metric, respectively (Fig. S8; Table S10). These values exceeded the variance explained when considering samples across all females (CLR transformed abundance matrix R^2^=0.0146, weighted UniFrac matrix R^2^=0.0030, and unweighted UniFrac matrix R^2^=0.0037 in Table S2). This increase suggests that while reproductive state has a consistent effect, its signal is masked in population-level analyses by the overwhelming inter-individual variation in gut microbial composition. By analyzing samples within single hosts, we partition out this inter-individual noise, revealing that the effect of reproductive state on microbial community structure is not just a population-level phenomenon but is consistently detectable within individual females.

Given this personalization, we tested if gut microbiota similarity changed between the same and different female baboon hosts and different reproductive states over time using temporal autocorrelation analyses. Pregnancy was the most personalized and dynamic phase of reproduction (orange lines in Fig. 4). Specifically, pairs of samples from pregnant females had the lowest Aitchison, weighted UniFrac, and unweighted UniFrac similarities of all reproductive states, whether those samples were from the same female and same pregnancy (Fig. 4A, 4D, 4G), the same female but a different pregnancy (except at the 6 reproductive-months lag for Aitchison similarity; Fig. 4B, 4E, 4H), or a different pregnant female baboon (Fig. 4C, 4F, 4I; Table S11). By contrast, ovarian cycling was the least personalized and dynamic phase of reproduction (green lines in Fig. 4). In all reproductive states, the highest gut microbial similarity occurred between pairs of samples collected from the same female host, in the same reproductive state, within 30 days of each other (left-hand sides of Fig. 4A, 4D, 4G). As expected, this similarity declined with increasing time between samples (Fig. 4A, 4D, 4G). In comparison, we observed lower similarity between samples from different reproductive events from the same female host (middle column of plots: Fig. 4B, 4E, 4H), and the lowest similarity between samples from different females in the same reproductive state, highlighting host-specific personalization (right-hand column of plots: Fig. 4C, 4F, 4I). In panel 4B, the slight increase in Aitchison similarity at the 6 reproductive-months lag may be an artifact of the reduced number of comparisons possible at this time lag (many samples can be collected within thirty reproductive days, but fewer samples can be collected between 150 and 180 reproductive days of each other, especially considering pregnancy lasts ∼180 days in total).

**Figure 4.**
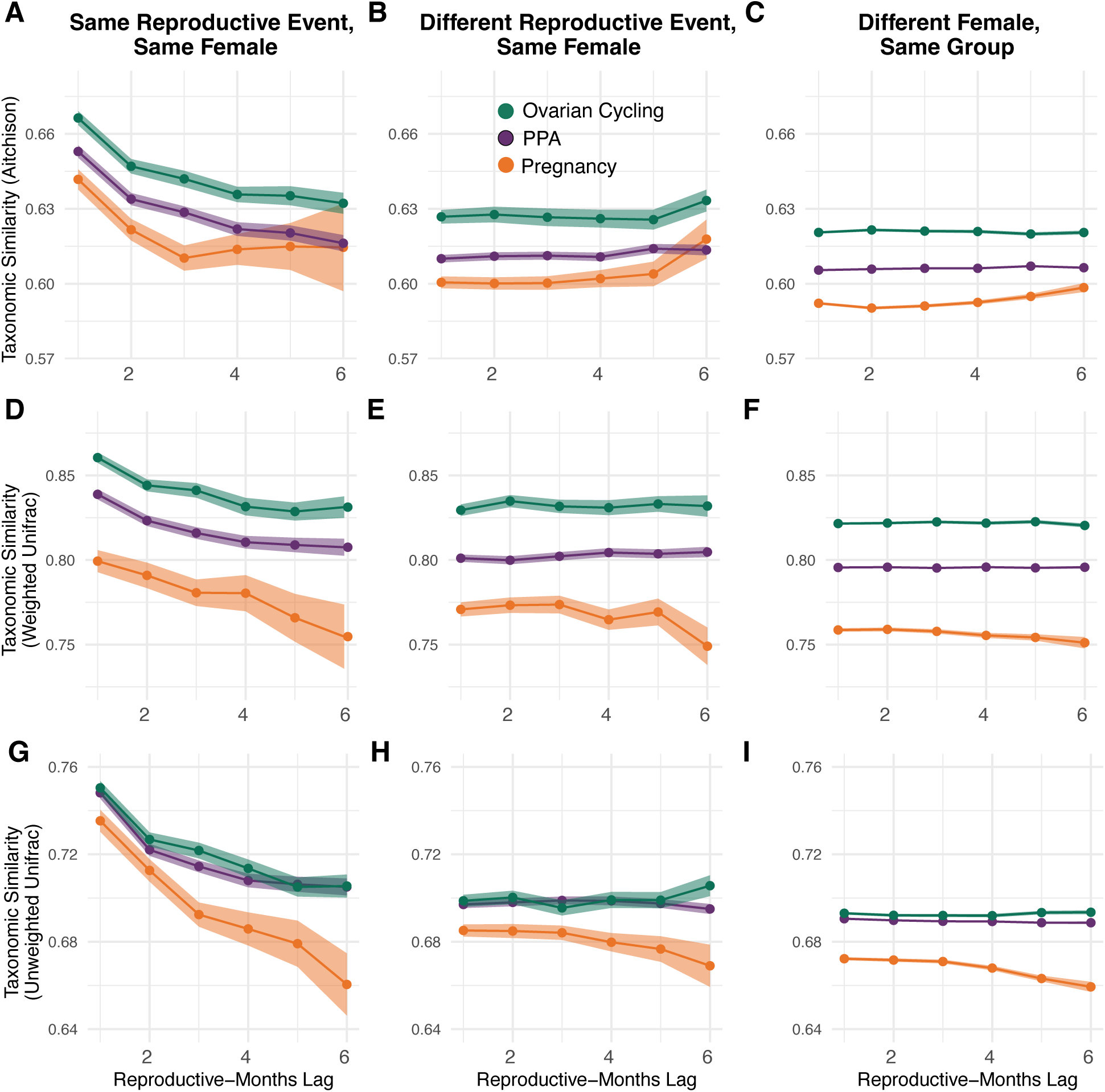
The pregnancy microbiota is more variable and personalized than other reproductive states. Plots show a temporal autocorrelation analysis of Aitchison similarities (A-C), Weighted UniFrac similarities (D-F), and unweighted UniFrac similarities (G-I) between pairs of gut microbiota samples from (A,D,G) the same reproductive event in the same female host, (B,E,H) different reproductive events in the same host, and (C,F,I) different hosts in the same social group. Points and lines are colored by reproductive state (green=ovarian cycling; purple=PPA; orange=pregnancy); bands around each line represent the 95% confidence interval. The x-axis shows how many months apart the samples were collected during the reproductive state in question (i.e., the reproductive-months lag). For instance, pairs of samples collected within 30 days of the same time point within a reproductive event are collected within a 1-month lag (the left-most data points on the x-axes); samples collected between 30 and 60 days of the same time point are collected within a 2-month lag (the second left most data points) and so on. Points higher on the y-axis have greater pairwise similarity, while points lower on the y-axis have lower pairwise similarity. All analyses include 4,124 samples from 115 females that had at least 10 total samples, including at least 2 samples from each reproductive state and only reproductive events with at least 2 samples. Statistical comparisons between points are in Table S11.

### 3. Fecal steroid hormone concentrations are associated with gut microbial composition, but the effects vary between reproductive states

Lastly, we tested how variation in steroid hormones—mean-centered fecal estrogen (fE), fecal progesterone (fP), and fecal glucocorticoid (fGC) concentrations—were associated with variation in gut microbiota, controlling for reproductive state and other covariates. We found that high fE and fP concentrations were significantly associated with high ASV richness, Shannon diversity, and phylogenetic diversity (Table S12; Fig. 5A-B; Fig. S9). However, these effects varied depending on reproductive state. For instance, the relationships of fE and fP with all three alpha diversity measures were weaker during pregnancy than during PPA (Table S12; Fig. 5, Fig. S9). In addition, the relationship between fE and alpha diversity was weaker during pregnancy than during ovarian cycling, and the relationship with fP and alpha diversity was weaker during ovarian cycling than PPA (Table S12). This difference is perhaps because fE and fP are already high during pregnancy, and are likewise elevated during ovarian cycling compared to PPA (Fig. 1A, Fig. 5A-B, Fig. S9A, B, D, E).

**Figure 5.**
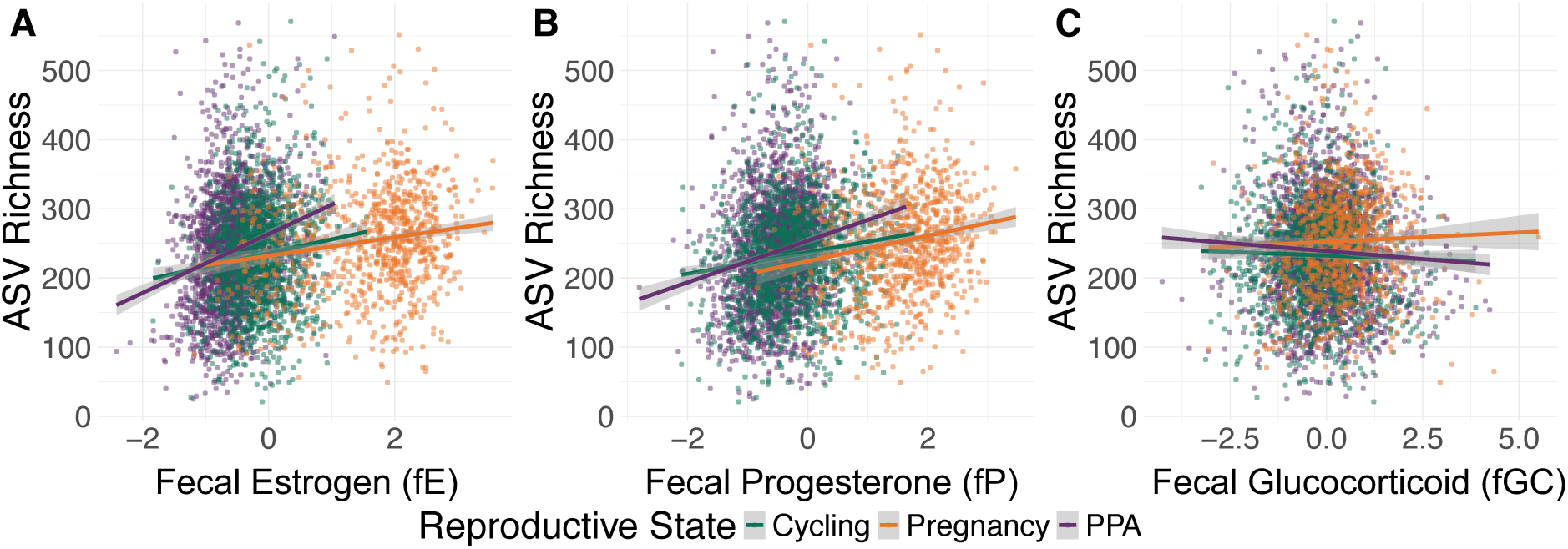
Fecal estrogen, progesterone, and glucocorticoids predict ASV richness. Plots show ASV richness as a function of the concentrations (ng/g) of (A) fE, (B) fP, and (C) fGC in that sample. Hormone values are corrected for the time to extraction and assay and mean-centered to 0 (see methods). Each point represents a fecal sample, colored by reproductive state, with linear model best fit line for cycling samples in green, pregnancy samples in orange, and PPA samples in purple. Model results are in Table S12.

In contrast, high fGC concentrations were significantly associated with lower alpha diversity (Table S12; Fig 5C; Fig. S9C, S9F). These results for fGC were largely consistent between reproductive states, with small significant interaction effect only found in two pairwise comparisons (pregnancy versus PPA: ASV richness β=4.73, p=0.021 and Shannon diversity β=0.046, p=0.049; Fig. 5C, Fig. S9C, S9F; Table S12). All three hormones were associated with statistically significant variation in beta diversity (i.e., between sample variation in microbiota composition), though these effects were very small (CLR abundance matrix R^2^ range=0.0022-0.0049, p=0.001; weighted UniFrac matrix R^2^ range=0.0024-.014, p≤0.013; unweighted UniFrac matrix R^2^ range=0.0024-0.0094, p≤0.014; Table S13). As with alpha diversity, we found small but significant interaction effects for beta diversity between reproductive state and fE (R^2^ range=0.0016-0.0030, p=0.001), fP (R^2^ range=0.0006-0.0030, p≤0.016), and fGC (R^2^ range =0.0007-0.0017, p≤0.002; Table S13).

In terms of individual microbial taxa, fE concentrations were associated with differential abundances of 126 ASVs (31.4% of 401 ASVs), 14 families, and one phylum, controlling for reproductive state and other covariates (Fig. 6A-C, 6G-H; Table S14-17). Likewise, fP concentrations were associated with differential abundances of 165 ASVs (41.1% of 401 ASVs), 31 families, and three phyla (Fig. 6D-F, 6I-J; Table S14-17). Again, many of the ASVs and families that were associated with sex steroid hormones were members of phylum Bacillota. However, the relationships of fE and fP to the abundances of individual taxa varied across reproductive states (Fig. 6). For instance, fE was associated with 93 ASVs during PPA, but only 27 ASVs during pregnancy and 53 during cycling, and fP was associated with 123 ASVs in PPA samples but only 84 ASVs in pregnancy and 24 ASVs in cycling samples (Fig. 6). Three families —Christensenellaceae, Monoglobaceae and Oscillospriales UCG-10—were positively associated with both fE and fP; all of these families belong to phylum Bacillota (Table S14-17). Seven families were negatively associated with fE and fP (Table S14-17). Four of these families were members of phylum Actinomycetota (Actinobacteria), specifically, Atopobiaceae 2, Coriobacteriaceae 2, an unidentified Coriobacteriales 2 family, and an unidentified Actinobacteria 2 family (Table S14-17). In addition, families Lactobacillaceae, Pasteurellaceae 2, and Brachyspiraceae were negatively associated with both fE and fP (Table S14-17). At the phylum level, Verrucomicrobiota (Tenericutes) was positively associated with fE during PPA and cycling and with fP during PPA (Table S14-17). Actinomycetota was negatively associated with fP during PPA and Thermodesulfobacteriota (Thermodesulfobacteria) was negatively associated with fP during pregnancy (Table S14-17).

**Figure 6.**
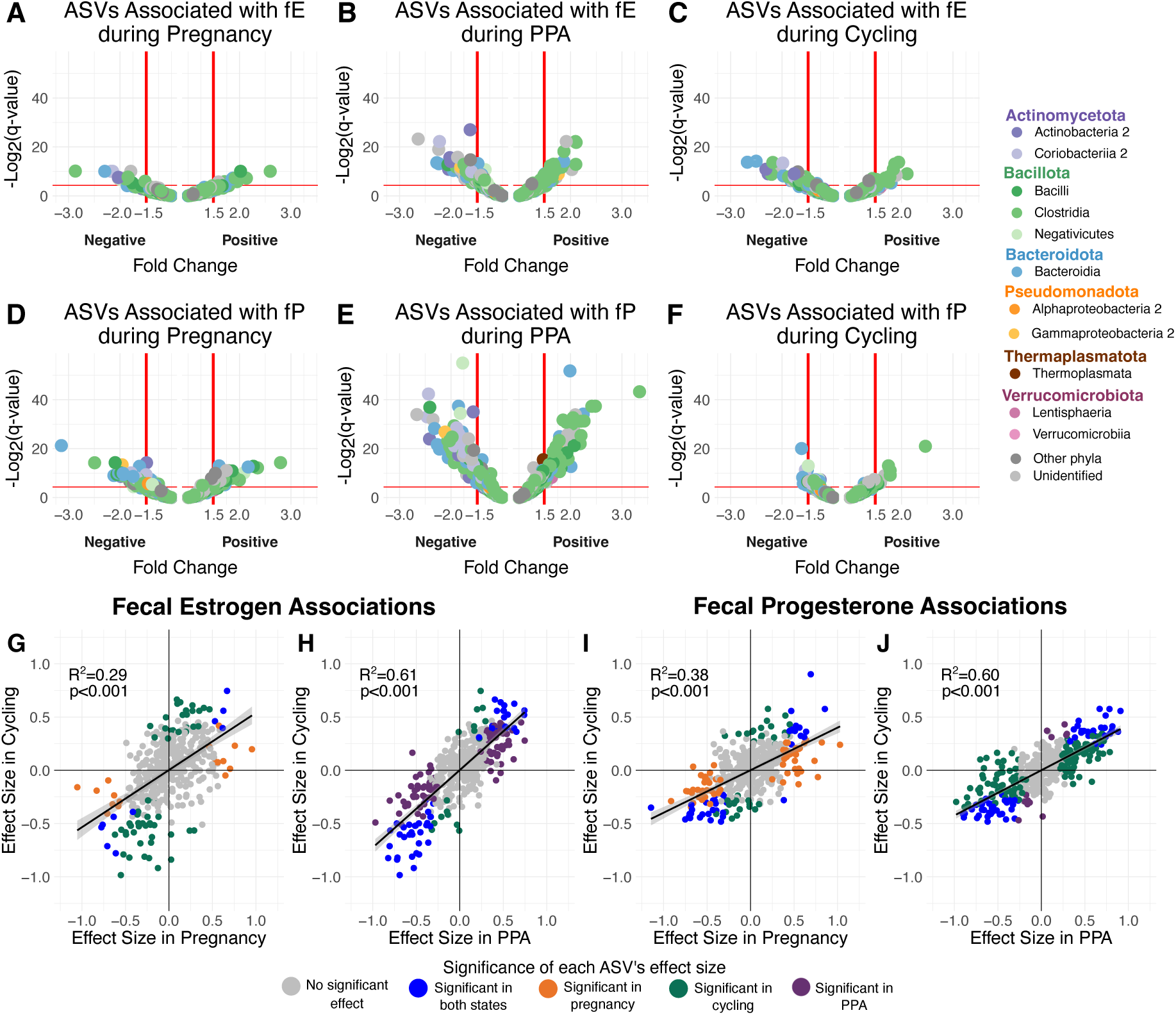
Estrogen and progesterone predict relative abundances of ASVs, with different effects in each reproductive state. (A, B, and C) Volcano plots showing the association between fE concentrations during pregnancy (A; Table S15), PPA (B; Table S16), and cycling (C; Table S17) and the abundances of the 401 ASVs present in 20% or more of samples. Each point represents an individual ASV and the color of each point represents the ASV’s assigned class. (D, E, and F) Volcano plots showing the association between fP concentrations during pregnancy (D; Table S15), PPA (E; Table S16), and cycling (F; Table S17) and the abundances of the 401 microbial ASVs. Each point represents an individual ASV and the color of each point represents the ASV’s assigned class. (G, H, I, and J) Scatterplots showing the correlation between effect sizes for individual ASV abundances with (G) fE in cycling and pregnant samples, (H) fE in cycling and PPA samples, (I) fP in cycling and pregnant samples, and (J) fP in cycling and PPA samples. Each point represents one of the 401 ASVs present in 20% or more of samples, colored by the reproductive states(s) where there is a significant (q<0.05) association between CLR-transformed relative abundance of that ASV and fE or fP. R^2^ and p-values are from linear models on the points, with linear model best fit lines shown on the plots.

In contrast to fE and fP, fGC concentrations predicted the relative abundances of relatively few taxa. Across all samples and within each reproductive state, fGC was strongly associated with only five ASVs (1.2% of 401 ASVs), one family, Gastranaerophilaceae, and no phyla (Table S14-17).

## DISCUSSION

The gut microbiota is connected to several aspects of female physiology that change during reproduction (e.g., immunity, metabolism)—yet scientific understanding of how and why the gut microbiota shapes and is shaped by these changes is still developing [3, 35, 120]. Leveraging a large, longitudinal dataset of samples from 169 female baboons across 14 years, we found strong, predictable changes in females’ gut microbiota as they move through cycling to conception, pregnancy, and lactation. These changes were evident both between reproductive states (cycling, pregnant, and postpartum amenorrhea) and between trimesters within pregnancies. While the gut microbiota changed relatively little during ovarian cycling, once females conceived, their microbiota shifted in microbial diversity and composition, especially during the 2^nd^ and 3^rd^ trimesters. This “pregnancy microbiota” was relatively personalized to individual females (compared to other reproductive states). The signature of pregnancy on the microbiota then faded in the first months of postpartum amenorrhea (PPA), so that as females in PPA approached cycling resumption, their microbiota returned to the composition they exhibited during ovarian cycling. Reproductive hormones were also important predictors of gut microbial composition, perhaps because these hormones affect microbial environments or because their production is directly influenced by certain gut microbes [121–123]. Below we discuss the implications of these results, focusing on the pregnancy microbiota, its personalization, and the connections between hormones and gut microbial composition. Together, our research provides a critical springboard to investigate the role of the gut microbiota in mammalian reproduction and its evolutionary consequences.

### A pregnancy microbiota in baboons

Across mammals, female gut microbiota are reshaped during pregnancy, with distinct effects on microbial taxonomic diversity and abundances [4, 5, 35, 124–126]. Such “pregnancy microbiota” have been identified and studied, mostly cross-sectionally, in humans, captive animals, and a handful of wild mammal populations [44, 61, 125, 127–137]. Here we document a pregnancy microbiota in wild baboons, largely driven by community shifts in mid- and late-pregnancy. We also confirm that the gut microbial signature of pregnancy is consistent both within and across individuals, and we identify specific windows during which the characteristic pregnancy microbiota develops.

Many taxa associated with pregnancy in our study, such as those in the phyla Pseudomonadota (Proteobacteria) and Bacillota (Firmicutes), have known functions related to pregnancy physiology in other mammals [130, 133–135, 138]. For instance, tight regulation of the immune system is an important aspect of pregnancy, with early and late pregnancy characterized as pro-inflammatory compared to the anti-inflammatory second trimester [21, 139, 140]. Consistent with this pattern, we found that the Pseudomonadota family Burkholderiaceae was more abundant during late pregnancy in baboons. A recent study found that a *Burkholderia* strain stimulates the immune system in an insect model [141]; our results motivate testing whether related microbial taxa interact with the immune system in pregnant mammals. Pseudomonadota as a whole are broadly implicated in host inflammation [142, 143] and are more abundant during pregnancy in baboons. Pregnancy-associated variation in the phylum Bacillota in our samples also may be related to immune function and/or host metabolism. Some Bacillota taxa promote weight gain through increased capacity for energy harvest and production of the short-chain fatty acid butyrate [144–146], which in turn has anti-inflammatory effects on the host [35, 147]. Other Bacillota taxa, including members of the family Lachnospiraceae that we identified as pregnancy-associated, such as members of the genera *Ruminococcus* and *Roseburia*, can induce a pro-inflammatory state via the upregulation of Treg and Th2 cells [3, 130, 144].

Understanding the relationship between pregnancy and alpha diversity is important because higher gut microbial diversity is often assumed to be healthier for both hosts and their microbiota [148, 149]. This idea is largely based on theoretical work predicting that microbial diversity produces greater functional redundancy and hence resilience in the face of perturbations [150–152]. More diverse microbiota might also be more challenging for pathogens to invade [153, 154]. However, the relationships between alpha diversity and pregnancy thus far reported in the literature vary between species, suggesting that biological variation in species’ physiology, phylogenetic histories, and/or environments at the time of sampling may be important in determining this relationship [155–157]. For instance, our observation that pregnant baboons have higher microbial alpha diversity during pregnancy contrasts with the growing consensus that humans have lower diversity during pregnancy compared to non-pregnancy and in later pregnancy compared to earlier pregnancy [3, 11, 61, 125, 144] (but see [158–160]). Similarly, among non-human primates, two studies identified lower alpha diversity during pregnancy [44, 133] and four found no difference in diversity as a function of reproductive state [131, 132, 161, 162]. No previous non-human primate research has found elevated alpha diversity during pregnancy as we observed in baboons. One distinctive feature of baboons is their year-round breeding, which is neither tightly seasonally-linked nor synchronized across females [51, 69]. Four of the six studies cited above are, in contrast, on highly seasonal breeders; non-seasonally breeding geladas and moderately seasonal white-faced capuchins were both found to have no differences in alpha diversity during pregnancy [132, 161]. Longitudinal sampling of other non- or weakly-seasonal breeders may be a fruitful step towards understanding these differences in alpha diversity between primate species. Our result, in combination with other studies of non-human animals that show considerable variation in patterns of pregnancy-associated gut microbial diversity [44, 128–133, 135–138, 161–170], emphasizes the complexity of the relationship between pregnancy and alpha diversity across mammals.

One likely source of this variability is the well-documented variation across mammals in patterns of maternal energetic investment during pregnancy, which is affected by the level of placental invasiveness, fetal brain and body size, the number of fetuses in a single pregnancy, and the length of gestation [171–175]. Primates exhibit a unique suite of pregnancy characteristics compared to other mammals of similar body size, including longer gestations, slower rates of fetal growth, and larger neonate bodies and brains [172, 176]. These traits allow pregnant primates, including humans, great flexibility in adjusting to the physiological demands of pregnancy, because the daily energetic requirements for fetal maintenance and development are relatively low [172, 176]. Studying other species with shorter gestations, faster rates of fetal growth, and/or smaller neonates will be essential for comparisons in a literature largely dominated by work on humans. A second source heterogeneity may be methodological: the timing of sample collection in cross-sectional studies of pregnancy is highly variable and often only includes a single time point. This sampling regime may contribute to the conflicting results in the literature. As more research investigates the gut microbiota and female reproduction in other mammals, comparative longitudinal studies will be a productive path to understanding what shapes the relationship between reproductive state and gut microbial alpha diversity across species.

### Gut microbial personalization during reproduction

We expected that the well-documented personalization of the gut microbiota [46, 63] would be least marked during pregnancy, based on the reasoning that the relatively consistent suite of physiological changes required to carry a fetus to term would lead to consistent microbial changes across females. Contrary to this expectation, we found that personalization was strongest during pregnancy, with gut microbiota being less similar between pregnancy samples—whether from the same or from different females—than during other states. Typical explanations for between-host microbiota differences, including host genotype, priority effects, and functional redundancy [13, 46, 151, 157, 177], likely contribute to some of this variation. Indeed, although these differences in within-reproductive state variability are small, they may reflect increased microbial volatility driven by shifts in immune regulation and host physiology during pregnancy that alter the strength and consistency of host-mediated selection, resulting in greater inter-individual and longitudinal variation [11, 30, 157, 178, 179]. Three additional processes may also contribute to personalization during pregnancy in particular.

First, pregnant females exhibit much greater variance in estrogen and progesterone levels than females in other reproductive states. Specifically, variance in fE is significantly higher during pregnancy than PPA (variance in pregnancy=0.838, SD=0.916; variance in PPA=0.182, SD=0.427; Levene’s test F=577.72, Bonferroni-corrected q<0.001); the same pattern occurs when we compare pregnancy to cycling (variance in cycling=0.205, SD=0.453; F=377.12, q<0.001). Variance in fP is also significantly higher during pregnancy than PPA (variance in pregnancy=0.502, SD=0.708; variance in PPA=0.232, SD=0.482; F=171.92, q<0.001), although not compared to cycling (variance in cycling=0.442, SD=0.665; F=3.62, q=0.17). Interestingly, while these hormones are connected to gut microbial composition in other states, fE and fP are less robust predictors of microbial diversity during pregnancy itself. Rather than a direct linear decoupling, the high variance in both hormones and microbiota may reflect a highly personalized physiology during pregnancy, where uniform predictive relationships break down. The pattern of greater variance during pregnancy could also hold true for other traits that influence reproductive physiology and vary across individuals and over the lifetime, including psychosocial stress, gene expression, and metabolic status [180–182], though we cannot explicitly test this prediction in our data set.

Second, the unique genotype of each fetus might also contribute to greater personalization during pregnancy. This possibility follows from the observation that fetal genotype can have direct impacts on maternal physiology, including stimulating maternal appetite [183, 184]. Third, interactions between fetal and maternal characteristics could contribute to the diversity of physiological responses to pregnancy [184]. For instance, in humans, hyperemesis gravidarum (severe nausea and vomiting during pregnancy) is caused by an increase in serum levels of hormone GDF15, which is largely produced by the feto-placental unit, not the mother herself [185]. However, pre-pregnancy maternal GDF15 levels, which have a genetic basis, modulate the risk of developing hyperemesis gravidarum even as pregnancy GDF15 levels vary [185]. Given the gut microbiota’s ties to immune, metabolic, and psychosocial health [21, 145, 179], we suspect that both fetal and maternal variation play a role in shaping personalized pregnancy-related microbial dynamics. While our study design does not allow us to explicitly test these ideas, they are a promising direction for future longitudinal work following multiple pregnancies across individual females.

### Estrogen and progesterone predict gut microbiota

Fluctuations in steroid hormones may be one factor driving reproduction-associated changes in gut microbial communities. Our results support this idea for the two sex steroid hormones we evaluated—estrogen and progesterone, which predict increases in gut microbial alpha diversity—but not for glucocorticoids. Empirical studies thus far have disagreed about the nature of the correlations between sex steroid hormone levels and gut microbial diversity: some studies have reported positive relationships, others have reported negative relationships, and still others have found no relationship [38, 122, 123, 186–192]. Conflicts in the literature may arise in part from the special nature of the populations under study (i.e., women with hormonal and metabolic disorders, menopausal women, or ovariectomized lab animals [122, 186, 187], instead of population-typic or natural levels), or from species-level differences. For example, in the one other study that focused on hormone levels and gut microbiota across reproduction in a natural primate population, progesterone was negatively correlated with gut microbial Shannon diversity, estrogen showed no relationship with gut microbial diversity, and alpha diversity was also lowest during pregnancy, when progesterone levels are highest [44]. All three patterns differ from our observations, highlighting the complexity of the reproductive hormone-microbiota interface. More longitudinal research across a larger array of species will be important to elucidate the extent and drivers (e.g. host phylogeny, physiology, behavior [35, 63, 155, 156]) of species differences in the relationships between reproductive hormones and the gut microbiota.

Nevertheless, our study does reveal internally consistent effects for several hormone-associated taxa with the potential to affect host health. For instance, we found an inverse relationship between fP and Pseudomonadota, which is consistent with the idea that progesterone suppresses host immunity [139, 193] (some Proteobacteria are pathogenic and pro-inflammatory [142, 143]). Matching patterns found in pregnant animals, Bacillota taxa are often positively associated with fE and fP. Clostridiaceae ASVs tend to have positive associations with both hormones, which is expected given that taxa in this family have β-glucuronidase genes, which enable bacteria to deconjugate biologically-inactive conjugated estrogen into its biologically active form, which is then reabsorbed by the host and returned to circulation [25, 122, 187]. Likewise, some species in this family can upregulate progesterone [39]. The Bacillota family Christensenellaceae and ASVs in this family are also positively associated with fE and fP. Butyrate, the SCFA mainly produced by Bacillota taxa, has direct effects on sex hormones, increasing their secretion from ovarian granulosa cells [40, 42, 147].

We found that the relationship between reproductive hormones and gut microbial community composition—especially alpha diversity—differed across reproductive states. Specifically, estrogen and progesterone were less robust predictors of alpha diversity during pregnancy than other reproductive states, a novel finding of this study. As previously noted, this weaker predictive power suggests that while there is greater hormonal variance during pregnancy, it does not predict a uniform microbial response. One explanation for this pattern is that both of these hormones are already high during pregnancy (Fig.1A), and the relationship between hormone levels and bacterial diversity may plateau at high hormone levels. Alternatively, other physiological changes that are unique to pregnancy, such as tight regulation of immunity and metabolism, may exert effects on the microbiota that outweigh those of hormones [35].

### Conclusions and future directions

Here we leveraged a unique, longitudinal data set from a wild mammal population to demonstrate that the gut microbiota is distinct during pregnancy and tied to host physiological shifts in pregnant baboons. We found some evidence for gut microbial personalization during pregnancy, which may be due to the effects of variation in, and interactions between, maternal and fetal traits that impact microbial composition. Hormonal fluctuations may be a mechanism by which the host directly influences the gut microbiota and the gut microbiota in turn influences host physiology. However, the relationship between hormones and the microbial community is not consistent across reproductive states and is likely modulated by other aspects of host physiology like immune and metabolic function. Fine-grained, regularly timed biological sample collection will enable further investigation of how the relationship between hormones and the gut microbiota is impacted by host physiology, and experimental approaches can additionally test causal mechanisms.

Moving forward, metagenomic sequencing techniques will be useful in elucidating what genes and microbial functions are present in gut microbes associated with reproductive states and hormones, rather than potentially present based on taxonomy. Although our 16S-based analyses can only measure taxonomic differences, our work provides strong motivation for future metagenomic studies. Longitudinal data sets with paired hormone and microbial measurements will be critical to advancing our understanding of these relationships, particularly in light of the personalization found in the gut microbiota during reproduction. In particular, despite a growing understanding of the importance of the gut microbiota during pregnancy, there is considerable disagreement across studies and species about how community diversity and taxonomic abundances shift during gestation. Longitudinal data collected from different mammalian species during pregnancy will help reveal if these differences between studies represent true biological differences due to host phylogeny, physiology, or ecology, or are artifacts of cross-sectional sampling. Further, future research connecting gut microbiota to fitness will help propel the field past descriptions of patterns and towards an improved evolutionary understanding of host-microbe associations.

## DECLARATIONS

### Ethics approval

This research was approved by the IACUCs at Duke University and the University of Notre Dame and the Ethics Council of the Max Planck Society. It adhered to all the laws and guidelines of Kenya. In Kenya, our research was approved by the Wildlife Research Training Institute (WRTI), Kenya Wildlife Service (KWS), the National Commission for Science, Technology, and Innovation (NACOSTI), and the National Environment Management Authority (NEMA).

### Consent for publication

Not applicable.

### Availability of data and materials

All data for these analyses are available on Dryad (https://doi.org/10.5061/dryad.5tb2rbphw). The 16S rRNA gene sequencing data are deposited on EBI-ENA (project ERP119849) and Qiita (study 12949). Code is available at the following

GitHub repository: https://github.com/CASouthworth/Southworthetal_GutMicrobiotaReproduction.

### Competing interests

The authors declare that they have no competing interests.

### Funding

We are grateful to the National Science Foundation (NSF) and the National Institutes of Health (NIH) for sustaining the ABRP’s multi-decadal research. We particularly acknowledge the following grants for the work described here: NSF DEB 1840223 (E.A.A., J.A.G.), NIH R01 AG071684 (E.A.A., S.C.A), NIH R21 AG055777 (E.A.A.), NIH R01 AG053330 (E.A.A.). We also acknowledge support from the NSF GRFP (C.A.S.), the Jack Kent Cooke Foundation Graduate Scholarship (C.A.S.), the P.E.O. Scholar Award (C.A.S.), the NSF Integrated Data Science Fellowship (L.B.), the Duke University Population Research Institute P2C-HD065563 (pilot award to J.T.), and Notre Dame’s Eck Institute for Global Health (E.A.A.) and Environmental Change Initiative (E.A.A.). Current support for field-based data collection also comes from the Max Planck Institute for Evolutionary Anthropology, and we thank Duke University, Princeton University, and the University of Notre Dame for financial and logistical support. The authors declare that the funding bodies listed here did not serve any role in the design of the study and collection, analysis, and interpretation of data and in writing the manuscript.

### Authors’ contributions

C.A.S. and E.A.A. designed the research. S.C.A., E.A.A., J.T., M.R.D., L.R.G., and J.A.G. produced the data. C.A.S. and L.B. analyzed the data. C.A.S. and E.A.A. wrote the manuscript with contributions from all authors.

## Acknowledgements.

We thank Jeanne Altmann for her instrumental role in stewarding the Amboseli Baboon Research Project. We also thank the University of Nairobi, the Kenya Institute of Primate Research (KIPRE), the National Museums of Kenya, the members of the Amboseli-Longido pastoralist communities, the Enduimet Wildlife Management Area, Ker & Downey Safaris, Air Kenya, and Safarilink for their cooperation and assistance in the field. Particular thanks go to the Amboseli Baboon Project long-term field team (R.S. Mututua, S. Sayialel, J.K. Warutere, I.L. Siodi, and L. Musembei), and to T. Wango and V. Oudu for their assistance in Nairobi. The baboon project database, Babase, is expertly managed by J. Gordon and C. Broderick. Database design and programming are provided by K. Pinc. For a complete set of acknowledgments of funding sources, logistical assistance, and data collection and management, please visit http://amboselibaboons.nd.edu/acknowledgements/.

## Notes

https://doi.org/10.5061/dryad.5tb2rbphw

https://github.com/CASouthworth/Southworthetal_GutMicrobiotaReproduction

### Competing Interest Statement

The authors have declared no competing interest.

### Summary of Updates

Revisions based on peer review comments.

