## Supplemental Methods & Figures for "Longitudinal changes in gut microbiota across reproductive states in wild baboons"

**Supplementary methods.**

*Fecal sample collection.* Feces were collected from known individual baboons within 15 minutes of defecation and preserved in 95% ethanol. Samples were stored in Amboseli at ambient temperature then transported to the University of Nairobi where they were freeze-dried, sifted, and stored at -20°C until transportation to the United States. Once in the US, samples were stored at -20°C until fecal powder was loaded into MoBio and QIAGEN PowerSoil kit for 96-well plates [1] (~0.05 g fecal powder per well). After sample loading, plates were sealed and stored at -80°C until DNA extraction.

*DNA sequencing.* To construct 16S rRNA gene sequencing libraries, polymerase chain reaction was used to amplify a ~390 bp-long fragment encompassing the V4 region [2, 3]. Amplicons were quantified via the Quant-iT PicoGreen dsDNA Assay Kit (ThermoFisher/Invitrogen cat. no. P11496) and equal amounts of amplicon DNA from each sample (70 ng) were pooled and cleaned using AMPure XP beads (Beckman Coulter). Libraries were sequenced on the Illumina HiSeq 2500 using the Rapid Run mode (2 lanes per run) and sequences were single indexed on the forward primer and 12 bp Golay barcoded to enable a high level of multiplexing [3].

*Analysis pipeline and quality filtering.* During demultiplexing, we used the *--create-fastq-for-index-reads* argument to prevent barcode misassignment, retained sequences with a maximum error estimate (maxEE) <0.1, and applied a minimum length=150 bases for Illumina library adapter removal and to remove too-short reads [4–6]. Read pairs were merged using DADA2 and we removed potential chimeric reads. Additional quality control steps in the original, 17,277 profile data set revealed no strong relationship between sample storage length and DNA concentration (b=2.0x10^-4^, p=0.064) [7]. Using sequencing technical replicates (182 replicates from 30 samples) across the original 40 plates, we also confirmed that technical replicates clustered together in a Bray-Curtis dissimilarity matrix rather than with their sequencing plate, indicating that true biological differences between samples were stronger than plate-based batch effects [7].

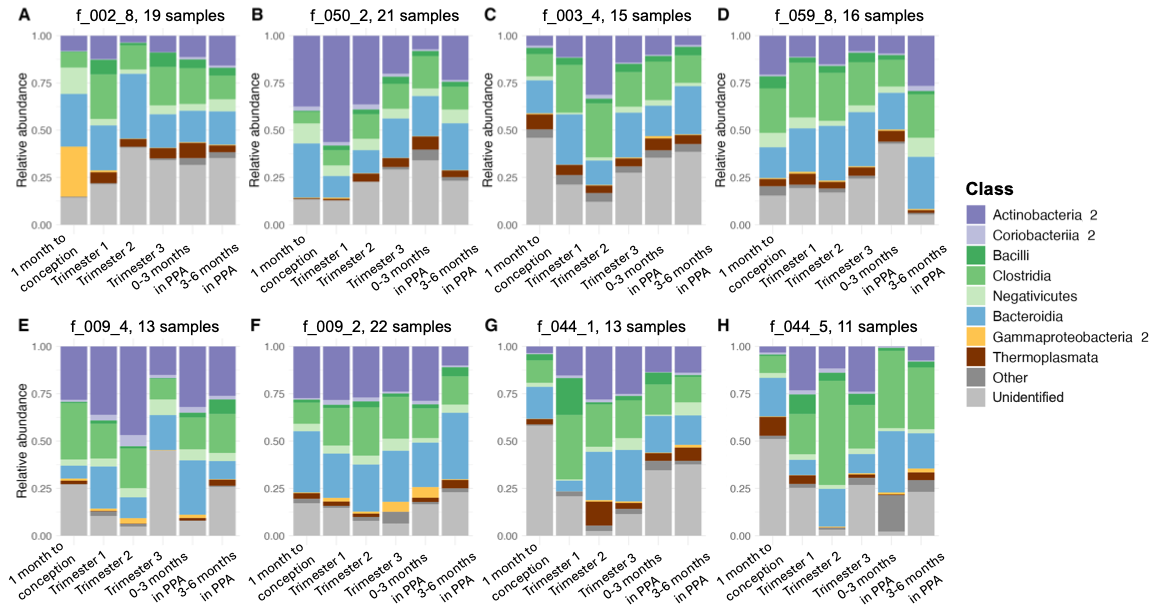

**Figure S1. Microbiota trajectories in individual female baboons across six phases of reproduction.** Average relative abundance of bacterial classes as females transition from the month before conception, trimester 1, trimester 2, trimester 3, 0 to 3 months postpartum, and 3 to 6 months postpartum). (A-D) each show one reproductive event for each of four female hosts (f_002, f_050, f_003, and f_059). (E-F) and (G-H) show two different reproductive events in two female hosts (f_009 and f_044). Each host is represented by a unique three number code and each reproductive event by a number that indicates parity. “Unidentified” taxa are those not identified to the class level and “Other” taxa aggregates rare taxa identified to the class level but present in <40% of samples.

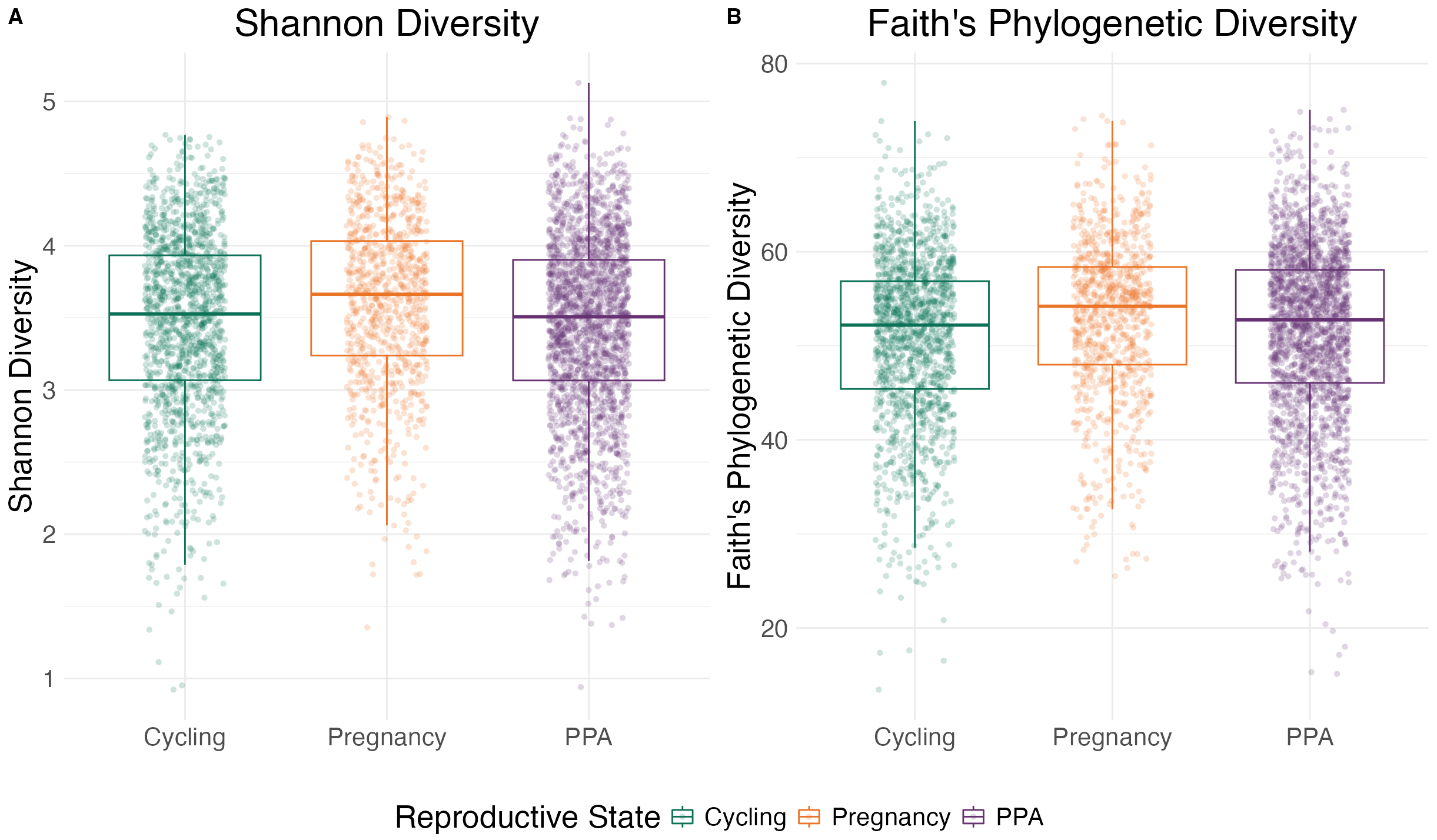

**Figure S2. Gut microbial alpha diversity is highest during pregnancy.** (A) Shannon diversity is higher during pregnancy than ovarian cycling (β=0.11, p<0.001) or PPA (β=0.14, p<0.001). (B) Faith’s phylogenetic diversity is higher during pregnancy than ovarian cycling (β=1.25, p<0.001) or PPA (β=0.97, p<0.001). Full model results are in Table S1.

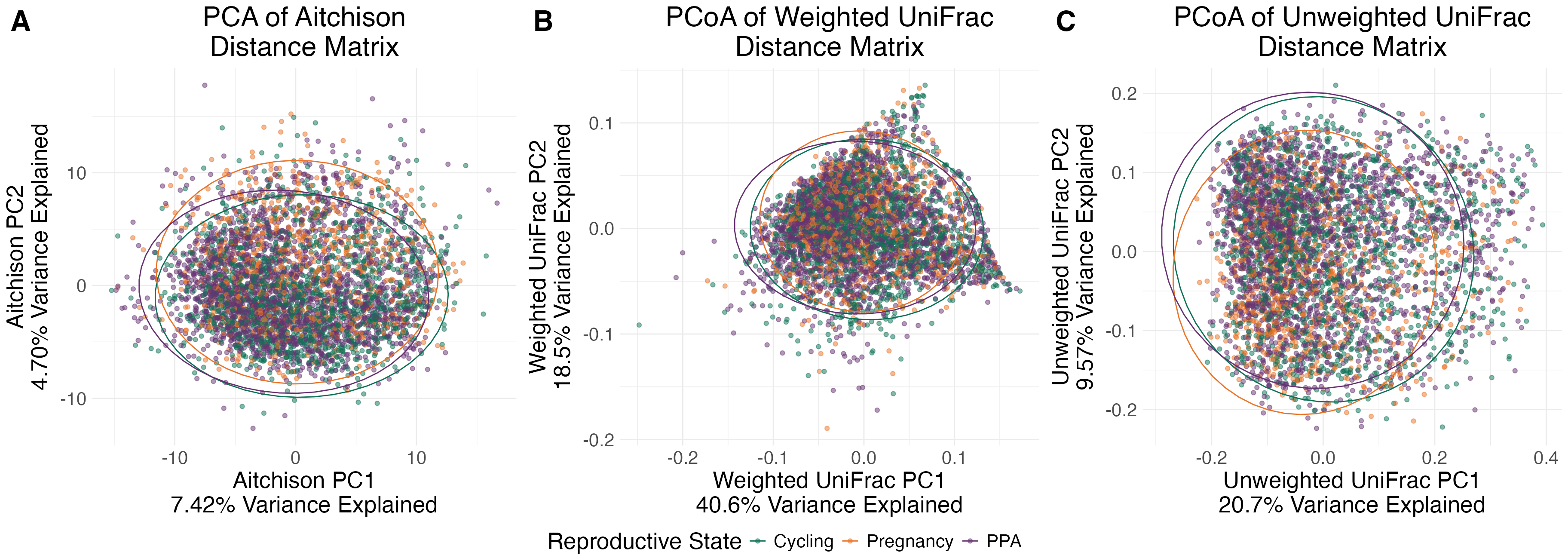

**Figure S3. Baboon gut microbiota do not cluster by reproductive state in ordination analysis.** Plots of the first two principal components (PCs) of an (A) Aitchison distance matrix, (B) weighted UniFrac distance matrix, and (C) unweighted UniFrac distance matrix. for all 4,462 samples. Each point is an individual fecal sample colored by reproductive state. Reproductive state explained 1.46% of the variation in the Aitchison distance matrix (ANOVA on RDA: R^2^ = 0.0146, p=0.001), 0.30**%** of the variation in the weighted UniFrac distance matrix (ANOVA on dbRDA: R^2^=0.0030, p=0.001), and 0.37% of the variation in the unweighted UniFrac distance matrix (ANOVA on dbRDA: R^2^=0.0037, p=0.001).

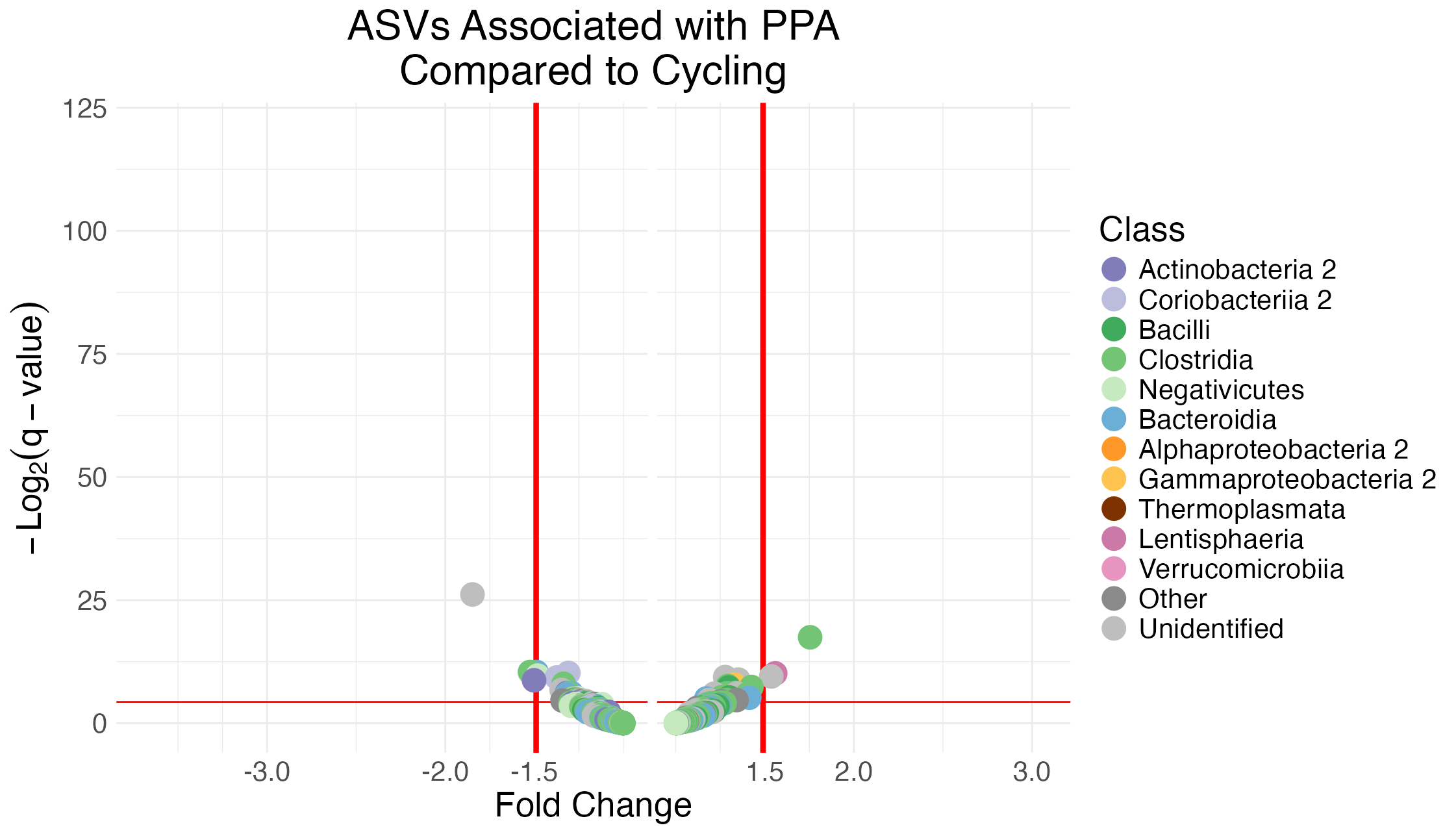

**Figure S4. Few gut microbial ASVs differ in abundance between PPA and ovarian cycling.** Volcano plot showing the effect of PPA compared to ovarian cycling on the abundances of the 401 ASVs present in at least 20% of samples. Each point represents an individual ASV and the color of each point represents the ASV’s assigned class. Points above the horizontal red line are statistically significant associations (q<0.05). Points to the left or right of the vertical red lines have relatively large fold changes of ≥1.5 or ≤-1.5. Model results used to generate this table are found in Table S4.

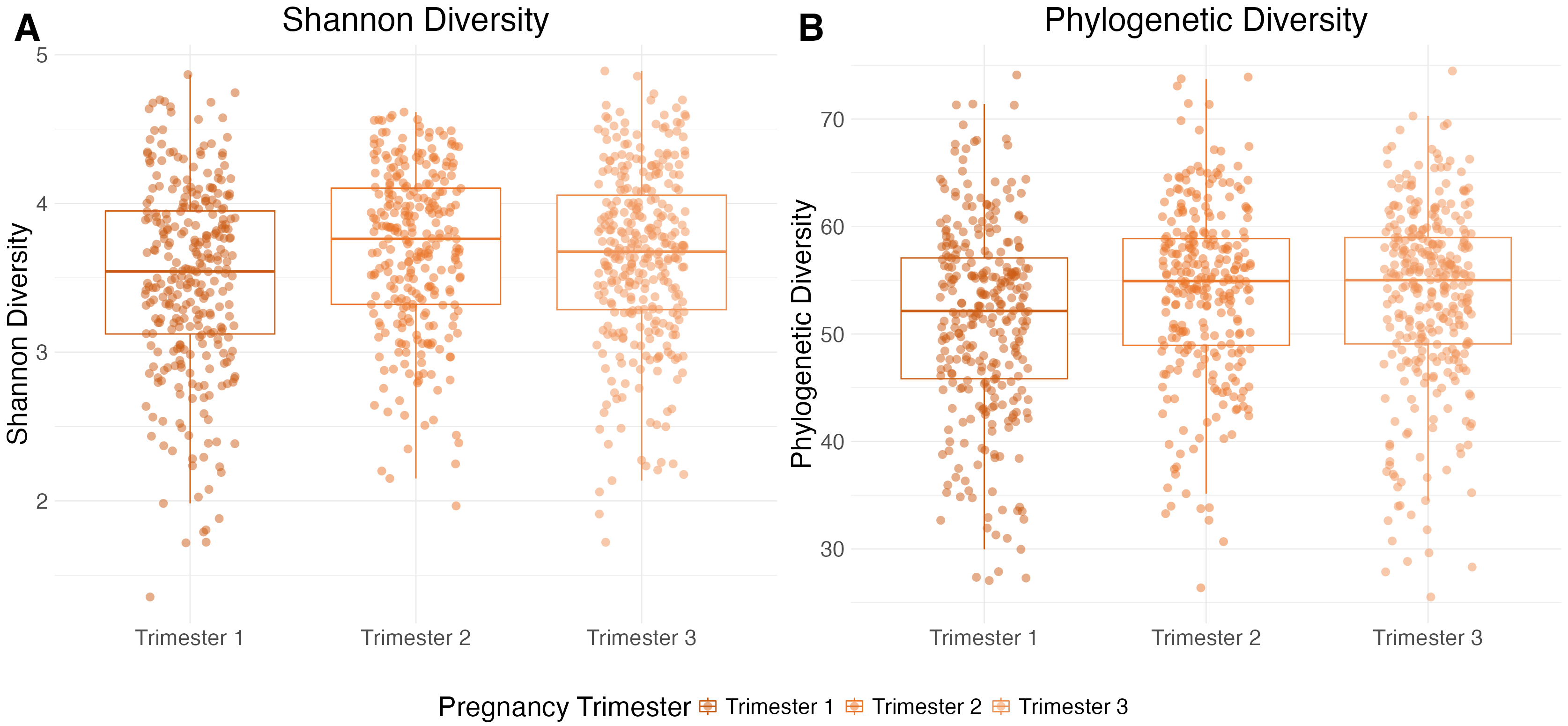

**Figure S5. Gut microbial alpha diversity is highest during trimesters 2 and 3 of pregnancy.** (A) Shannon diversity is higher during trimester 2 (β=0.21, p<0.001) and trimester 3 (β=0.15, p<0.001) compared to trimester 1. Trimesters 2 and 3 do not differ in Shannon diversity (β=-0.058, p=0.17). (B) Faith’s phylogenetic diversity is higher during trimester 2 (β=2.34, p<0.001) and trimester 3 (β=1.45, p=0.005) compared to trimester 1. Trimesters 2 and 3 do not differ in phylogenetic diversity (β=-0.89, p=0.085). Full model results are in Table S5.

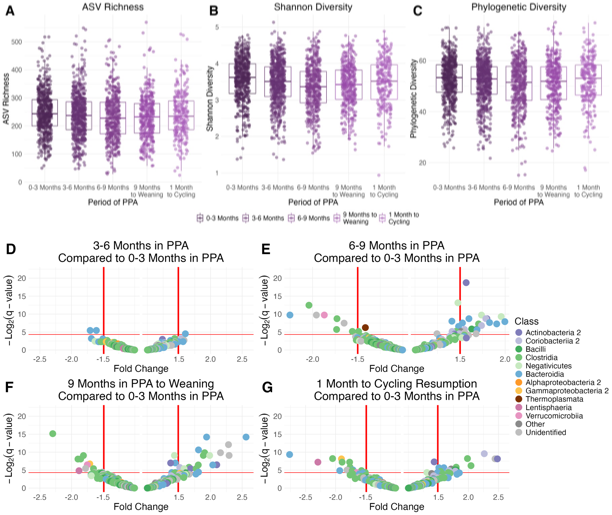

**Figure S6. Gut microbiota in the first three months of PPA are distinct from later PPA.** (A) ASV richness, (B) Shannon diversity, and (C) phylogenetic diversity as a function of the period of PPA. Alpha diversity is significantly higher during early PPA than later periods of PPA (Table S5). (D-G) Volcano plots showing the effect of three to six months in PPA (D), six to nine months in PPA (E), nine months in PPA to infant weaning (F), and one month to cycling resumption (G) compared to zero to three months in PPA on the abundances of the 401 bacterial ASVs. Each point represents an individual ASV and the color of each point represents the ASV’s assigned class. Points above the horizontal red line are statistically significant (q<0.05). Points to the left or right of the vertical red lines have fold changes of ≥1.5 or ≤-1.5. Model results used to generate panels D-G are in Table S8.

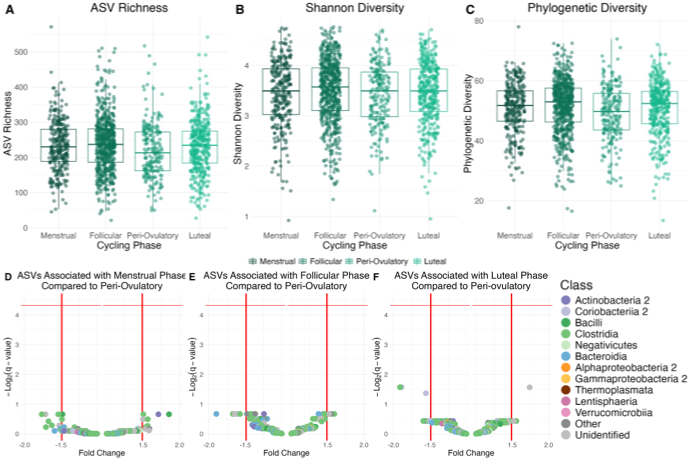

**Figure S7. Gut microbial changes are small across ovarian cycle phases.** (A) ASV richness, (B) Shannon diversity, and (C) Faith’s phylogenetic diversity as a function of ovarian cycle phase. Alpha diversity does not significantly differ between phases (Table S5). (D-F) Volcano plots showing the effect of the menstrual (D), follicular (E), and luteal (F) phases compared to the peri-ovulatory phase on the abundances of the 401 bacterial ASVs. Each point represents an individual ASV and the color of each point represents the ASV’s assigned class. Points above the horizontal red line are statistically significant (q<0.05). Points to the left or right of the vertical red lines have fold changes of ≥1.5 or ≤-1.5. Model results used to generate panels D-F are in Table S9.

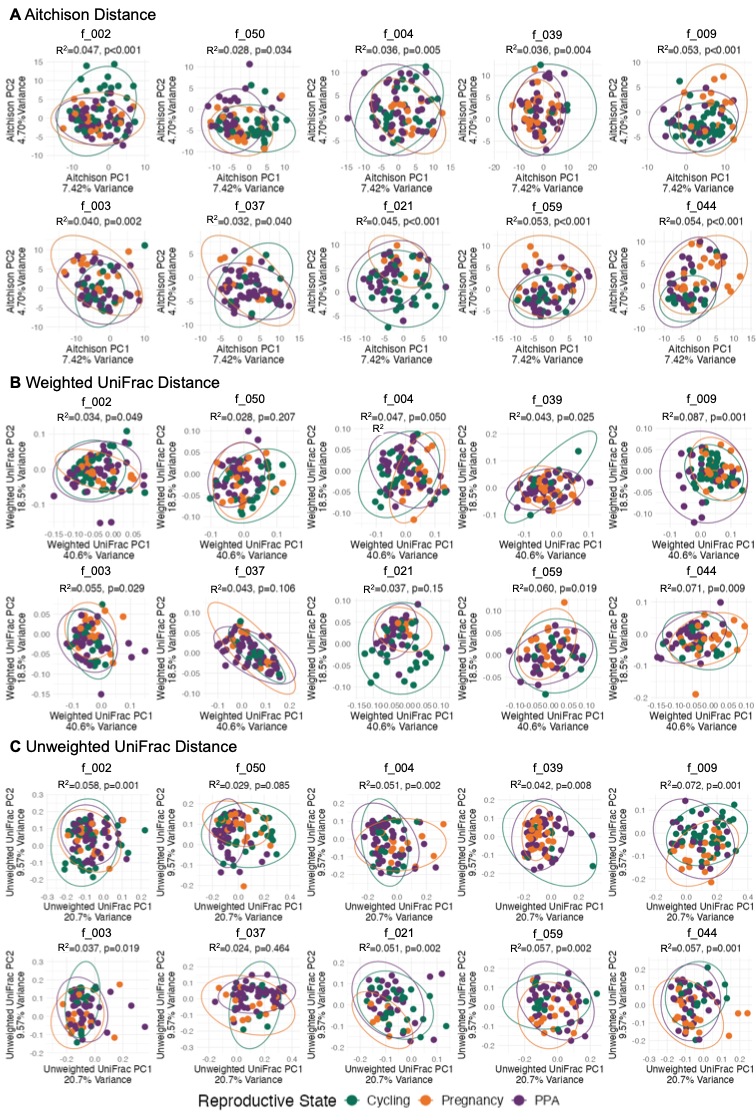

**Figure S8. Reproductive state explains more variation in microbiota composition within individuals, compared to across all samples.** Each plot shows the first two principal components of (A) Aitchison distances, (B) weighted UniFrac distances, and (C) unweighted UniFrac distances between samples within an individual female (female ID is indicated by a unique code above each plot). Each point represents one sample and points are colored by reproductive state. R^2^ and p-values are from ANOVAs on the RDAs (Aitchison distances) or dbRDAs (weighted and unweighted UniFrac distances) for each female in Table S10.

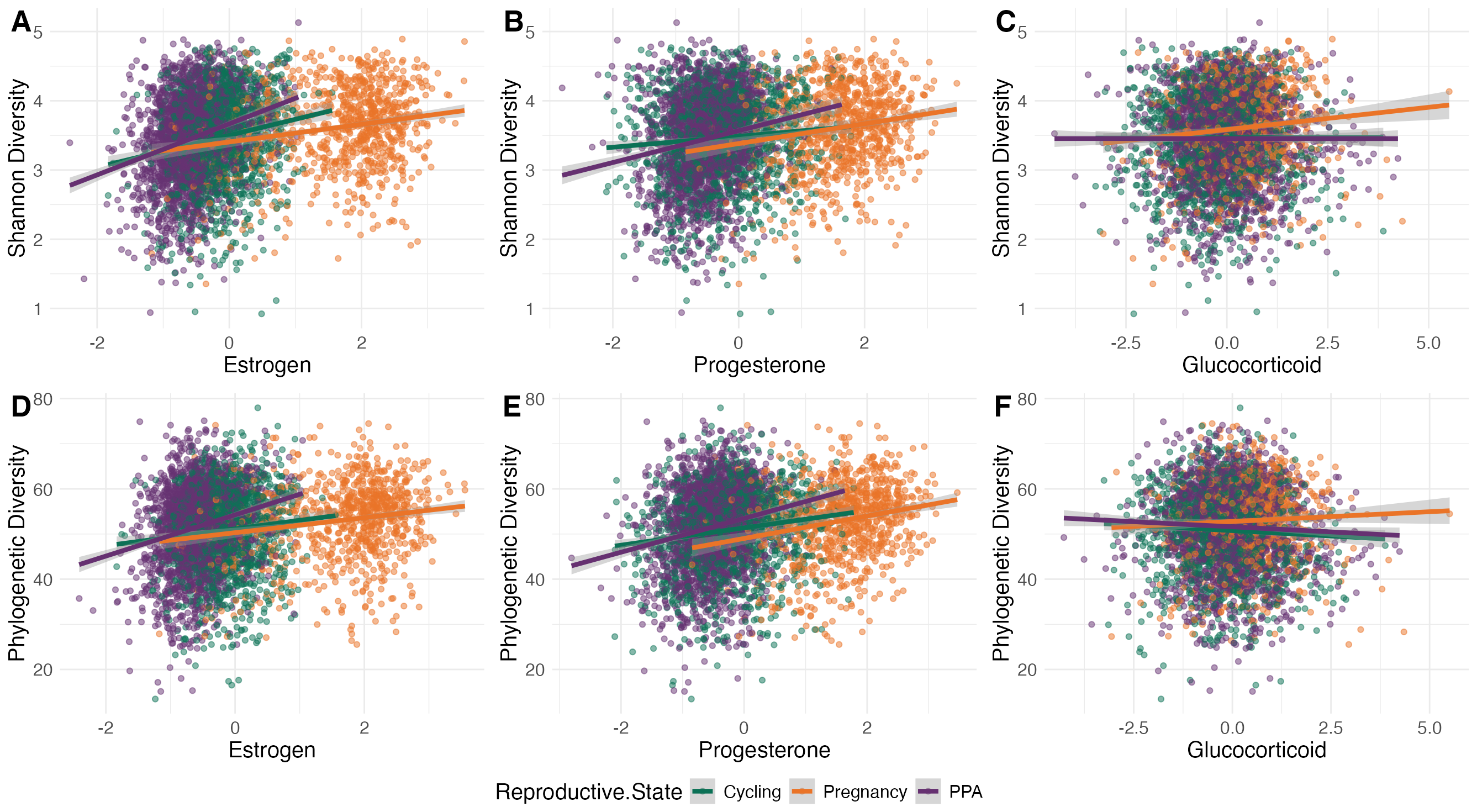

**Figure S9. Steroid hormones predict Shannon diversity and Faith’s phylogenetic diversity.** Plots show Shannon diversity in fecal samples as a function of the concentration of (A) fecal estrogen metabolites (ng/g), (B) fecal progesterone metabolites (ng/g), and (C) fecal glucocorticoid metabolites (ng/g) in that sample, and phylogenetic diversity as a function of (D) estrogen metabolites, (E) progesterone metabolites, and (F) glucocorticoid metabolites. Hormone values are corrected for the time to extraction and assay and mean-centered to 0 (see methods). Each point represents a fecal sample, colored by reproductive state, with linear model best fit line for cycling samples in green, pregnancy samples in orange, and PPA samples in purple. Model results are in Table S12.

**References.**

1. QIAGEN. MO BIO’s PowerSoil-htp 96 well soil DNA kit handbook. 2014.

2. Caporaso JG, Lauber CL, Costello EK, Berg-Lyons D, Gonzalez A, Stombaugh J, et al. Moving pictures of the human microbiome. Genome Biol. 2011;12:R50. https://doi.org/10.1186/gb-2011-12-5-r50.

3. Caporaso JG, Lauber CL, Walters WA, Berg-Lyons D, Huntley J, Fierer N, et al. Ultra-high-throughput microbial community analysis on the Illumina HiSeq and MiSeq platforms. ISME J. 2012;6:1621–4. https://doi.org/10.1038/ismej.2012.8.

4. Callahan BJ, McMurdie PJ, Rosen MJ, Han AW, Johnson AJA, Holmes SP. DADA2: High-resolution sample inference from Illumina amplicon data. Nat Methods. 2016;13:581–3. https://doi.org/10.1038/nmeth.3869.

5. Illumina. bcl2fastq2 conversion software v2.20. 2019.

6. Martin M. Cutadapt removes adapter sequences from high-throughput sequencing reads. EMBnet.journal. 2011;17:10–2. https://doi.org/https://doi.org/10.14806/ej.17.1.200.

7. Grieneisen L, Dasari M, Gould TJ, Björk JR, Grenier J-C, Yotova V, et al. Gut microbiome heritability is nearly universal but environmentally contingent. Science. 2021;373:181–6. https://doi.org/10.1126/science.aba5483.
